# Genomic and phenotypic characterisation of *Penicillium melinii*, a plant growth-promoting root endophyte that remains effective under salt stress

**DOI:** 10.64898/2026.11.16.766994

**Authors:** Sandra González-Sáyer, Laura Gutiérrez-Manso, Iván Devesa-Aranguren, Francesco Blasio, Lorena B. Conchillo, Carlos M. Conesa, Lydia Ugena, José Nolasco, Ana Vázquez-Mora, Soledad Sacristán, Elena Caro, Juan Carlos del Pozo, Javier Cabrera

## Abstract

*Penicillium melinii* ‘isolate 2’ is a root endophyte of *Arabidopsis thaliana* that promotes the growth of different plant species, but the genetic basis of this activity remains unknown. Here we combined Oxford Nanopore long reads and Illumina short reads to obtain a high-quality, near-chromosome-level assembly of 29.39 Mb in six nuclear scaffolds, two of them telomere-capped at both ends (97.6% complete BUSCOs), together with, to our knowledge, the first complete mitochondrial genome reported for the species (26,730 bp). Annotation predicted 9,566 protein-coding genes. Compared with two other plant growth-promoting *Penicillium* species, *P. bilaiae* and *P. citrinum*, the isolate encodes fewer carbohydrate-active enzymes, biosynthetic gene clusters and candidate effectors, pointing to an overall reduction of the secreted repertoire. *In vitro*, the isolate grew significantly above the control on 63 of 95 carbon sources, mainly carbohydrates, and was positive for 7 of 11 plant growth-promoting traits: siderophore production, phosphate and zinc solubilisation, ammonia release, and urease, proteolytic and lipolytic activities. Radial growth was unaffected by NaCl up to 150 mM and persisted at 1 M, whereas conidiation was more salt-sensitive. On medium containing 100 mM NaCl, inoculation increased *Arabidopsis* shoot fresh weight and root system growth. Genome mining identified candidate genes for these traits. Together, these results provide a genomic and phenotypic basis for evaluating *P. melinii* ‘isolate 2’ as an inoculant for saline soils and identify candidate determinants for functional validation.

## Introduction

Agricultural systems face escalating pressures that compromise both productivity and environmental stability. Intensification has driven progressive losses of soil organic matter, structural degradation, erosion and a decline in soil biota (Tilman et al., 2002; Shokri et al., 2025), while poor fertilizer use efficiency results in much of the nitrogen and phosphorus applied being lost to watercourses and the atmosphere, fuelling eutrophication and greenhouse gas emissions (You et al., 2023; Ludemann et al., 2024). Salt accumulation in agricultural soils adds a further dimension to this burden: salinisation ranks among the most severe forms of soil degradation and currently affects some 1.38 billion hectares, slightly over ten per cent of global land area, with a further billion at risk (FAO, 2024). Together, these pressures have renewed interest in sustainable alternatives to conventional agrochemical inputs.

Microbial inoculants have emerged as a promising complement to synthetic fertilisers and crop protection products. Under European Union Regulation (EU) 2019/1009, plant biostimulants are defined as products that stimulate plant nutrition processes independently of their own nutrient content, improving nutrient use efficiency, abiotic stress tolerance, quality traits and the availability of confined soil nutrients (European Parliament and Council, 2019; du Jardin, 2015). In a global meta-analysis of field trials, inoculation increased crop yield by an average of 16%, with the greatest response in dry climates, and improved nitrogen and phosphorus use efficiency (Schütz et al., 2018). Critically, plant growth promotion by microbial inoculants is consistently greater under salinity stress than in its absence (Tufail et al., 2021), which makes strains isolated from salt-affected environments particularly valuable: they combine inherent halotolerance with a demonstrated capacity to promote host plant growth under conditions where agrochemical inputs are least effective. The potential for widespread adoption is illustrated by the Brazilian soybean sector, where approximately 78% of the cultivated area, some 36.5 million hectares, is inoculated annually (Santos et al., 2019).

All plants in natural ecosystems appear to be symbiotic with fungal endophytes, a taxonomically diverse group that can profoundly benefit host fitness (Rodriguez et al., 2009). The mechanisms through which fungal isolates have been described to promote growth and alleviate stress fall into a limited number of classes. Hormonal modulation contains some of the best-supported mechanisms available in fungi. For auxin, the indole-3-pyruvate route is the biosynthetic pathway with the most extensive genetic validation in fungi (Reineke et al., 2008; Sardar and Kempken, 2018), but the link to the host is harder to establish. In *Ustilago maydis*, mutants blocked in this pathway reduced the IAA rise in infected tissue without compromising tumour formation (Reineke et al., 2008). Similarly, in *Metarhizium robertsii*, which promotes lateral root growth in Arabidopsis partly through an auxin-dependent mechanism, deleting the tryptamine-pathway gene Mrtdc reduced fungal IAA production, but not enough to affect seedling growth promotion (Liao et al., 2017). ACC deaminase provides the cleanest case: RNAi silencing of *Tas-acdS* in *Trichoderma asperellum* reduces its promotion of canola root elongation without altering root colonisation, and expressing the gene in *Escherichia coli* confers that promotion (Viterbo et al., 2010), although the chain linking the enzyme to stress ethylene was established in bacteria (Glick, 2014). Gibberellins have been quantified in *Penicillium* spp. culture filtrate (Khan et al., 2011a; Khan et al., 2013b) but to our knowledge no fungal gene has been manipulated to test the attribution. A second class of mechanisms are related to nutrient mobilisation. Phosphorus transfer has been demonstrated in a root endophyte by isotopic labelling, since *Colletotrichum tofieldiae* delivers ³³P to *Arabidopsis* shoots (Hiruma et al., 2016); mono- and dihydroxamate siderophores from *P. chrysogenum* improve the iron nutrition of cucumber and maize (Hördt et al., 2000); entomopathogenic fungi do so in sorghum on calcareous substrate, though the mechanism operating in planta remains unclear (Raya-Díaz et al., 2017); and insect-derived nitrogen transfer by Metarhizium is established by ¹⁵N labelling and is modulated by fungal ammonium permeases, whose deletion increased rhizoplane colonisation and nitrogen transfer (Behie et al., 2012; Moonjely et al., 2019).

The best-established agronomic application for the genus *Penicillium* concerns phosphorus: *P. bilaiae* is marketed as a phosphate-solubilising inoculant (Leggett et al., 2007), and the oxalic and citric acids responsible have been identified as the main acids produced directly (Cunningham and Kuiack, 1992). Beyond phosphorus, endophytic *Penicillium* species act through a broad set of mechanisms and across a range of stresses. *P. funiculosum* LHL06 secretes gibberellins that reprogramme soybean growth under copper stress (Khan et al., 2013a), *P. roqueforti* improves nutrient uptake in wheat on heavy metal-contaminated soils (Ikram et al., 2018), and a drought-tolerant *P. citrinum* from wheat leaves combines auxin and gibberellin production with phosphate and zinc solubilisation and siderophore secretion (Kaur and Saxena, 2023). Several isolates also promote host growth under salinity, including *P. minioluteum* LHL09 and *P. funiculosum* LHL06 in soybean (Khan et al., 2011a, b), *P. janthinellum* LK5 in tomato (Khan et al., 2013b), *P. chrysogenum* CM022 in wheat (Dargiri et al., 2025) and *P. olsonii* from the rhizosphere of *Aeluropus littoralis* (Tarroum et al., 2022).

Beyond these mechanisms, an inoculant must itself stay active in stressed rhizospheres such as those of saline soils. *Serendipita indica* lowers sodium in *Arabidopsis* tissue and expresses its ENA-type Na⁺-ATPases during co-cultivation under salinity (Lanza et al., 2019; Haro et al., 2023). Fungal salt-tolerance determinants (Na⁺-ATPases, Na⁺(K⁺)/H⁺ antiporters, K⁺ transporters and the high-osmolarity glycerol, HOG, pathway) are well established in yeast, to which genetic evidence for the antiporters and K⁺ transporters remains largely restricted (Ko and Gaber, 1991; Bañuelos et al., 1998), whereas phenotypes in filamentous fungi are partial or divergent. *Aspergillus nidulans* EnaA is required for growth at alkaline pH rather than for Na⁺ detoxification (Markina-Iñarrairaegui et al., 2020) and, unlike the osmosensitive *gpd1* mutants of *Saccharomyces* (Albertyn et al., 1994), its *gfdA* mutant still synthesises glycerol, the osmotic route depending instead on GldB (Fillinger et al., 2001; de Vries et al., 2003). HOG controls glycerol and arabitol accumulation in *Fusarium graminearum* (Zheng et al., 2012), whereas in *Beauveria bassiana* and *Magnaporthe* the HOG-regulated solute appears to be arabitol (Zhang et al., 2009; Dixon et al., 1999). Several *Penicillium* sp. isolates have been described to promote host growth under salinity conditions, including *P. minioluteum* LHL09 and *P. funiculosum* LHL06 in soybean (Khan et al., 2011a, b), *P. janthinellum* LK5 in tomato (Khan et al., 2013b), *P. chrysogenum* CM022 in wheat (Dargiri et al., 2025) and *P. olsonii* from the rhizosphere of *Aeluropus littoralis* (Tarroum et al., 2022). Persisting in the soil is not enough: the fungus must also colonise the root and be tolerated by host immunity. *Trichoderma virens* strains lacking the cerato-platanin Sm1 colonised maize roots normally but no longer protected the plants against *Colletotrichum graminicola* (Djonović et al., 2007). LysM effectors such as Ecp6 sequester chitin to prevent chitin-triggered immunity in pathogens (Bolton et al., 2008; de Jonge et al., 2010); in beneficial fungi, RiSLM is required for full mycorrhizal colonisation of *Medicago truncatula* (Zeng et al., 2020) and Tal6 protects *Trichoderma atroviride* hyphae and modulates plant defence (Romero-Contreras et al., 2019). Arabidopsis root endophytes retain large repertoires of plant cell wall-degrading enzymes, and the strains that colonise roots most aggressively tend to be detrimental to the host, a link attributed to the pectate lyase family PL1_7, whose overexpression increased root colonisation at the expense of plant performance (Mesny et al., 2021).

The mechanisms described above, share a common methodological constraint: they were characterised through targeted biochemical and genetic approaches that examine one pathway at a time, and results are frequently species-specific and sometimes contradictory. Genome-wide analysis addresses this constraint directly. Comparative genomics of mycorrhizal fungi revealed that the mycorrhizal habit has evolved independently on several occasions, through convergent losses of plant cell wall-degrading enzymes and lineage-specific suites of symbiosis-induced genes, a pattern invisible to single-gene surveys (Kohler et al., 2015). Similarly, profiling 24 *Penicillium* sp. genomes uncovered a secondary metabolite biosynthetic potential far exceeding what targeted biochemical characterisation had captured (Nielsen et al., 2017), and analysis of root-associated fungi linked specific effector and CAZyme repertoires to the endophytic lifestyle (Mesny et al., 2021). Rather than testing candidates one by one, genome-wide annotation provides a simultaneous, unbiased inventory of all gene families encoding growth-promoting and stress-tolerance functions. Comparing genomes across related species further separates machinery conserved across the genus from innovations restricted to individual lineages, allowing predictions about whether a given trait is likely to be shared or species-specific (Kohler et al. 2015, Petersen et al. 2023). For the genus *Penicillium*, genomic resources have expanded rapidly (Nielsen et al., 2017), yet most PGP-active species lack the high-quality, near-chromosome-level assemblies needed to fully resolve gene content, repetitive regions and secondary metabolite cluster architecture.

*Penicillium melinii* is a common soil fungus, reported among the most abundant species of boreal mixed-wood forests in Quebec (De Bellis et al., 2007), and it was one of six plant growth-promoting fungi recovered from the rhizosphere of the halophyte grass *Aeluropus littoralis* (Tarroum et al., 2021), where its mycelium increased the biomass of tobacco seedlings and its cell-free culture filtrate increased both root length and biomass. Independently, the strain designated ‘isolate 2’ was recovered from roots of a wild *Arabidopsis thaliana* specimen from central Spain as part of an endophyte collection (García et al., 2013). It has since been characterised as a beneficial root endophyte: it colonises the epidermis and cortex without invading the vascular cylinder, causes no symptoms, and promotes root development in *Arabidopsis in vitro* and in tomato and quinoa in the greenhouse, and increased tomato biomass and yield in the field, with the capacity conserved across strains of independent geographical origin (Gutiérrez-Manso et al., 2026). A short-read draft genome of the isolate has been reported (305 contigs ≥1 kb; Gutiérrez-Manso et al., 2026), but its fragmentation limits the analysis of gene content and genome organisation and the fungal genetic basis of the growth-promoting capacity has not been examined. A contiguous, fully annotated genome assembly is therefore the prerequisite for a systematic investigation of the molecular determinants underpinning its growth-promoting activity.

Here we report a high-quality near-chromosome-level genome assembly of the beneficial root endophyte ‘isolate 2’, obtained by combining Oxford Nanopore long reads and Illumina short reads, which supersedes the short-read draft of the same isolate (Gutiérrez-Manso et al., 2026), together with its mitochondrial genome. After confirming its phylogenetic position, we compare its predicted proteome with those of two other plant growth-promoting *Penicillium* species, *P. bilaiae* and *P. citrinum*, to ask whether its gene repertoire is distinctive or typical of the genus. We then examine the traits that may underlie its growth-promoting activity, combining carbon source profiling, *in vitro* assays of nutrient-mobilising and hydrolytic activities, and a genome-wide search for candidate genes. Finally, because the species has also been isolated from the rhizosphere of a halophyte whereas ‘isolate 2’ comes from a non-saline soil, we test whether the isolate tolerates salt and whether it promotes plant growth under salinity.

## Methods

### Fungal strain

The fungal isolate used in this study was *Penicillium melinii* ‘isolate 2’, previously characterised as a plant growth-promoting fungus by Gutiérrez-Manso et al. (2026) and deposited at the Spanish Type Culture Collection (CECT, Spain) under the terms of the Budapest Treaty, with deposit number CECT 21242. For experiments, the fungus was maintained on potato dextrose agar (PDA) plates for 7 days at room temperature. Spores were harvested by adding 1 mL water to each plate, and the suspension was adjusted to 10^9^ spores mL^-1^. A 10 µL aliquot of this suspension was used to inoculate each assay medium except where indicated otherwise. Unless stated otherwise, plates were incubated in a growth chamber (Conviron CMP6010) at 23 °C under a long-day photoperiod (16 h light / 8 h dark).

### Genomic analysis

#### High molecular weight DNA extraction

For DNA extraction, the strain was reactivated from glycerol stock on potato dextrose agar (PDA; Difco, BD Biosciences) and grown for 4 d at room temperature under ambient laboratory light. Three colonies were then transferred to a flask containing approximately 200 mL of autoclaved liquid medium (20 g/L yeast extract and 20 g/L glucose in Milli-Q water), which was incubated at 28 °C with orbital shaking at 200 rpm for 5 d. Mycelium was harvested by filtration through filter paper, collected in a 50 mL Falcon tube, frozen in liquid nitrogen, stored at −80 °C and subsequently freeze-dried in a LyoQuest freeze-dryer (Telstar, Spain) for 2 days. High-molecular-weight (HMW) DNA was extracted from 250 mg of fungal biomass following a protocol adapted from Apirajkamol et al. (2024). Biomass was pulverised under liquid nitrogen in three successive rounds, lyophilised and lysed in 600 µL of lysis buffer (50 mM Tris-HCl pH 8.5; 50 mM EDTA; 5% SDS; 1% 2-mercaptoethanol) with 20 µL of proteinase K (20 mg/mL) at 65 °C for 60 min with gentle inversion. After removal of cell debris, proteins were precipitated with 0.5 volumes of 3 M sodium acetate (pH 5.2) and DNA was precipitated with an equal volume of isopropanol, washed with 1 mL of 70% ethanol, air-dried and resuspended in 20 µL of TE buffer. All centrifugation steps were performed at 19,000 × g for 10–15 min. DNA concentration and purity were assessed with a NanoDrop One spectrophotometer (Thermo Scientific). Integrity was verified by agarose gel electrophoresis and by capillary electrophoresis on a TapeStation 4150 (Agilent) using the Genomic DNA ScreenTape assay.

#### Genome sequencing

This step was performed at Novogene. Briefly, a ligation library was prepared from HMW DNA of *P. melinii* ‘isolate 2’ using the Ligation Sequencing Kit SQK-LSK114 (Oxford Nanopore Technologies) following the manufacturer’s instructions, and sequenced on a PromethION FLO-PRO114M (R10.4.1) flow cell. Basecalling was performed with MinKNOW v26.01.15 (Oxford Nanopore Technologies (ONT), 2026b) using the high-accuracy (HAC) model. ONT reads were subsampled to 80× target coverage using Rasusa v3.0.0 (Hall, 2022; Furqon et al., 2026). Illumina paired-end short-reads from Gutiérrez-Manso et al. (2026) were quality-trimmed with fastp v1.0.1 (Chen et al., 2018), with automatic adapter detection for paired-end data, single-base trimming of the 5′ and 3′ ends of each read, and removal of reads in which more than 15% of bases were uncalled.

#### *De novo* assembly

Four assemblies were compared: Flye v2.9.6-b1802 (Kolmogorov et al., 2019) (--nano-hq --genome-size 30m), Raven v1.8.3 (Vaser and Šikić, 2021), Miniasm v0.3-r179 (Li, 2016) (preceded by all-versus-all overlap computation with minimap2 v2.28-r1209 (Li, 2018), -x ava-ont), and a hybrid assembly generated with MaSuRCA v4.1.4 (Zimin et al., 2013) combining the ONT and Illumina read sets. All assemblies were evaluated with QUAST v5.3.0 (Gurevich et al., 2013) and BUSCO v5.8.3 (Manni et al., 2021) (ascomycota_odb12). The MaSuRCA assembly was retained for downstream analyses.

The mitochondrial genome was assembled using a targeted read-extraction strategy. Reads mapping to three candidate mitochondrial scaffolds identified in the initial assembly were extracted with minimap2 v2.28 (Li, 2018), yielding 13,469 reads (94.8 Mb). Reads were downsampled to approximately 150× target coverage with Rasusa v3.0.0 (Hall, 2022), using a conservative size estimate based on the combined length of the three candidate scaffolds (∼56 kb), and assembled with Flye v2.9.5 (Kolmogorov et al., 2019) using the --nano-hq preset and a genome-size estimate of 56 kb, yielding a single circular contig of 26,601 bp (304× coverage). Circularity was independently confirmed with Circlator v1.5.5 (Hunt et al., 2015) using default nucmer-based end-overlap detection, and the linear start of the contig was set with Circlator fixstart at the ATG start codon of a Prodigal-predicted open reading frame at position 12875 of the pre-rotation assembly (Prodigal v2.6.3; Hyatt et al., 2010).. Final polishing with v2.2.2 (ONT, 2026a) against the full unsubsampled read set produced a 26,730 bp consensus. To further validate circularity, the polished consensus was duplicated in tandem, reads were remapped with minimap2 (-ax map-ont), and both terminal coverage uniformity and reads spanning the midpoint junction were examined.

#### Scaffolding, polishing and validation

The selected assembly was scaffolded with RagTag v2.1.0 (Alonge et al., 2022) using, as sequential references, the genome assemblies of the two most closely related species with publicly available sequences, *Penicillium atrosanguineum* (GCA_028827265.1) and *P. maclennaniae* (GCA_028827695.1), as identified by Gutiérrez-Manso et al. (2026). Divergence from both scaffolding references was quantified with MUMmer v4.0.0rc1 (Marçais et al., 2018) (dnadiff). Because these references belong to different species of the same genus, all gaps in the scaffolded assembly were assessed for long-read support: ONT reads were mapped with minimap2 v2.28-r1209 (-x map-ont), alignments were sorted with SAMtools v1.22.1 (Danecek et al., 2021), and primary alignments (MAPQ ≥ 20) spanning each gap with at least 1 kb of anchoring sequence on both flanks were retained. Of nine gaps identified, two intra-contig gaps had spanning read support (14 and 58 alignments, respectively); the two inter-scaffold junctions lacked spanning reads, and their orientation therefore rests on synteny with the reference assemblies. One round of short-read polishing was performed with Pilon v1.24 (Walker et al., 2014) using preprocessed Illumina reads mapped to the assembly with Bowtie2 v2.5.5 (Langmead and Salzberg, 2012) and coordinate-sorted with SAMtools v1.22.1. A subset of the raw ONT signal data was re-basecalled with Dorado v0.7.2 (ONT, 2024) (model: dna_r10.4.1_e8.2_400bps_hac@v5.0.0) and used for long-read scaffolding with ntLink v1.3.11 (Coombe et al., 2023) (k=32, w=100, z=1000). Scaffolds were screened for contamination with BlobTools v1.1.1 (Laetsch and Blaxter, 2017). Telomeric repeats were identified in the nuclear genome assembly using tidk v0.2.41 (Brown et al., 2025); the repeat unit was determined empirically with tidk explore over a 5–12 bp search range, and the genome was subsequently scanned for this motif.

#### Gene prediction and functional annotation

Structural gene prediction was performed using Funannotate v1.8.17 (Palmer and Stajich, 2020). The assembly was repeat-masked prior to gene calling using RepeatMasker v4.2.2 (Smit et al., 2013–2015), as implemented in funannotate mask. As no RNA-seq data are available for *P. melinii*, transcript evidence was derived from publicly available RNA-seq reads of *P. janthinellum* (NCBI BioProject PRJNA779195), the most closely related *Penicillium* species with public transcriptome data. Reads were aligned to the soft-masked assembly with HISAT2 v2.2.1 (Kim et al., 2019) (--dta, --score-min L,-1,-0.6, --mp 4,2, --rdg 2,1, --rfg 2,1, --pen-noncansplice 4, --max-intronlen 10000, -k 5), and alignments were coordinate-sorted with SAMtools v1.22.1. A custom non-redundant *Penicillium* protein database was constructed from NCBI reference proteomes of the nine *Penicillium* species most closely related to *P. melinii* ‘isolate 2’ (Gutiérrez-Manso et al., 2026; Supplementary Table 1) and clustered at 90% sequence identity with CD-HIT v4.8.1 (Fu et al., 2012). To identify the annotation strategy yielding the most complete gene models, four strategies were evaluated within the Funannotate predict pipeline: (i) ab initio prediction without external evidence; (ii) the NCBI NR protein database only; (iii) *P. janthinellum* RNA-seq alignments combined with the NCBI NR protein database; and (iv) *P. janthinellum* RNA-seq alignments combined with the *Penicillium* species-specific protein database. Annotation completeness for each strategy was assessed using BUSCO v5.8.3 (ascomycota_odb12, n = 2,826; Manni et al., 2021) to select the final gene models.

Functional annotation of predicted proteins was performed with InterProScan v5.59-91.0 (Jones et al., 2014) against the Pfam-A 35.0 (Mistry et al., 2021), SUPERFAMILY 1.75 (de Lima Morais et al., 2011) and SMART 7.1 (Letunic et al., 2012) databases with GO term and pathway annotation, and with eggNOG-mapper v2.1.13 (Cantalapiedra et al., 2021) against eggNOG DB v5.0.2 (Huerta-Cepas et al., 2019). Mitochondrial genes were annotated with MITOS2 v2.1.10 (Donath et al., 2019) using the RefSeq 89 fungal reference dataset, genetic code 4, and the --best option. To identify potential candidate genes across the whole proteome, we used a large language model (Claude Opus 5.5, Anthropic) to query the annotation table for genes that could contribute to plant growth-promoting functions, based on the functional traits obtained in our analyses. All candidates named in the text were reviewed by the authors, who verified each assignment against the published literature.

#### Secretome, CAZyme and BGC annotation

The predicted proteome was screened to identify putative secreted proteins. Signal peptides were predicted independently with SignalP v6.0h (Teufel et al., 2022) and Phobius v1.01 (Käll et al., 2004); only proteins for which both tools predicted a signal peptide and Phobius predicted no transmembrane helices were retained as secretome candidates. Subcellular localisation of the remaining candidates was further assessed with DeepLoc v0.1 (Almagro Armenteros et al., 2017); only proteins classified as extracellular were retained as the candidate secretome. Potential effector proteins within the candidate secretome were identified using EffectorP v3.0 (Sperschneider and Dodds, 2022). Protease family assignments were made using the MEROPS database v12.5 (Rawlings et al., 2018), as implemented in Funannotate annotate.

Carbohydrate-active enzymes (CAZymes) were annotated by searching the predicted proteome against the dbCAN HMM database V10 (Zheng et al., 2023) using hmmscan from the HMMER v3.4 package (Eddy, 2011); hits were filtered to an i-Evalue of ≤ 1×10⁻¹⁵ and a domain coverage of ≥ 0.35, and overlapping or redundant family and subfamily hits on the same locus were collapsed to the single highest-scoring call. Secondary metabolite biosynthetic gene clusters were predicted with antiSMASH v8.0 (Blin et al., 2025) using the web server in relaxed detection mode. Functional annotations for secretome candidates were retrieved from the funannotate genome annotation by gene identifier.

#### Phylogenomic placement

Phylogenomic placement of *P. melinii* ‘isolate 2’ among *Penicillium* was inferred using single-copy BUSCO orthologues. Sixteen reference assemblies were retrieved from NCBI GenBank and analysed together with that of *P. melinii* ‘isolate 2’: plant-growth-promoting species (*P. bilaiae*, *P. citrinum*, *P. olsonii*, *P. oxalicum*, *P. janthinellum*, *P. chrysogenum*), postharvest pathogen references (*P. digitatum*, *P. italicum*, *P. expansum*), additional *Penicillium* diversity (*P. corylophilum*, *P. atrosanguineum*, *P. maclennaniae*, *P. canescens*, *P. griseofulvum*, *P. rubens*) and *Aspergillus fumigatus* as outgroup. BUSCO v6.0.0 (Manni et al., 2021) was run in genome mode against the eurotiales_odb10 lineage dataset for each assembly. Single-copy orthologues present in all 17 taxa (completeness threshold 100%; -psc 100.0) were aligned with MAFFT v7.526 (Katoh and Standley, 2013) (--auto), trimmed with trimAl v1.5.rev0 (Capella-Gutiérrez et al., 2009) (-automated1), and concatenated into a supermatrix of 2,223,900 amino acid positions using BUSCO_phylogenomics (McGowan, 2022). A maximum-likelihood phylogeny was inferred from the concatenated alignment using FastTree v2.2.0 (Price et al., 2010), with branch support estimated using SH-like local support values.

#### Comparative genomic analysis

Comparative genomic analyses were conducted among *P. melinii* ‘isolate 2’, *P. bilaiae* ATCC 20851 and *P. citrinum* B9. These species were selected on the basis of experimentally confirmed plant growth-promoting activity (Leggett et al., 2007; Chen et al., 2023) and the availability of predicted proteomes. The predicted proteome of *P. bilaiae* ATCC 20851 (v1.0) was obtained from the JGI MycoCosm database. The predicted proteome of *P. citrinum* B9 was obtained from NCBI GenBank (assembly accession GCA_036320845.1; Lau et al., 2024). Both proteomes were annotated using the same functional annotation pipeline applied to *P. melinii* ‘isolate 2’ to enable direct comparison.

Functional annotation profiles were compared across the three proteomes using COG category assignments from eggNOG-mapper v2.1.13 (Cantalapiedra et al., 2021) and domain annotations from InterProScan v5.59-91.0 (Jones et al., 2014). Secretome compositions were compared among the three species using candidate secretomes predicted for each proteome following the pipeline described above. Orthologous groups were inferred with OrthoFinder v3.1.2 (Emms and Kelly, 2019) under default parameters applied to the three predicted proteomes; orthogroups present in all three species were classified as core, and those exclusive to *P. melinii* ‘isolate 2’ were classified as species-specific. Genes within species-specific orthogroups were cross-referenced against the functional annotation, secretome classification, CAZyme annotation, and EffectorP v3.0 predictions generated for *P. melinii* ‘isolate 2’ as described above. EffectorP-positive candidates within species-specific orthogroups were further characterised by sequence similarity searches against the NCBI non-redundant protein database using blastp (BLAST+ v2.17.0+; Camacho et al., 2009; accessed September 2026), retaining up to ten target sequences per query.

### Phenotypic characterisation

#### Enzymatic activity

Lipase, protease, cellulase, laccase and amylase activities were screened qualitatively on selective solid media, based on the appearance of a halo or a colour change around the colony.

Lipase activity was tested on peptone agar (10 g peptone, 5 g NaCl, 0.1 g CaCl_2_·2H_2_O, 16 g agar, 1 L distilled water, pH 6 supplemented with 1 mL of 1% Tween 20 solution to each 300 mL medium), autoclaved separately and added once the base medium had cooled, to avoid surfactant degradation during co-sterilisation. Plates were incubated for 14 days; a zone of precipitation/opacity (calcium salt crystals) around the colony was recorded as positive for lipase activity (Hankin and Anagnostakis, 1975; Sierra, 1957).

Protease activity was evaluated on glucose–yeast extract–peptone (GYP) agar (1 g glucose, 0.1 g yeast extract, 0.5 g peptone, 16 g agar, 1 L Milli-Q water, pH 6 supplemented with 0.4% gelatine), autoclaved separately and mixed with the base medium before pouring. After 14 days, plates were flooded with saturated ammonium sulphate solution (76 g (NH_4_)_2_SO_4_ in 100 mL Milli-Q water) for 1 h; a clear halo around the colony indicated proteolytic activity (Hankin and Anagnostakis, 1975).

Cellulase activity was assessed on yeast extract–peptone agar (0.1 g yeast extract, 0.5 g peptone, 16 g agar, 1 L distilled water) amended with 0.5% carboxymethylcellulose (CMC). After 10 days, plates were flooded with 0.2% aqueous Congo red for 15 min and destained with 1 M NaCl for 15 min; an unstained halo around the colony indicated cellulolytic activity (Teather and Wood, 1982).

Laccase activity was tested on GYP agar amended with 0.005% 1-naphthol. After 14 days, a colour change from colourless to blue around the colony was recorded as positive for laccase activity (Sunitha et al., 2013).

Amylase activity was tested on GYP agar amended with 2% soluble starch. After 14 days, plates were flooded with iodine solution (1% I_2_ in 2% KI); a clear halo around the colony indicated amylase activity (Hankin and Anagnostakis, 1975).

#### Urease and ammonia production

Urease production was tested on Christensen’s medium (1 g peptone, 1 g glucose, 5 g NaCl, 2 g KH_2_PO_4_, 15 g agar, 0.012 g phenol red, 1 L distilled water) supplemented with filter-sterilised (0.22 µm) urea (0.5 mL of a 20% solution per 4.5 mL medium). Plates were inoculated with the spore suspension and incubated for 14 days; a colour change from yellow to pink was recorded as positive for urease activity (Christensen, 1946).

Ammonia production was assessed in peptone water (10 g L^-1^ peptone, 5 g L^-1^ NaCl, pH 7), following Cappuccino and Sherman (1992). Tubes containing 10 mL peptone water were inoculated with 10 µL of the spore suspension and incubated at 28 °C for 72 h with shaking (180 rpm) in the dark; an uninoculated tube served as a negative control. After incubation, 1 mL of Nessler’s reagent was added to each tube; development of a yellow-to-brownish-orange colour was recorded as positive for ammonia production.

#### Phosphate solubilisation

Phosphate-solubilising ability was tested on Pikovskaya (PVK) agar containing (g L^-1^): 10 glucose, 5 Ca_3_(PO_4_)_2_, 0.5 yeast extract, 0.5 (NH_4_)_2_SO_4_, 0.2 NaCl, 0.2 KCl, 0.1 MgSO_4_·7H_2_O, 0.002 MnSO_4_·H_2_O, 0.002 FeSO_4_·7H_2_O and 15 agar. Tricalcium phosphate was added before autoclaving and the medium was kept homogeneous while pouring, as the insoluble salt renders the plates opaque. Three medium variants were assayed: the standard, unadjusted PVK medium (pH ≈ 5.7), and PVK adjusted to pH 7 and pH 8 by adding previously autoclaved 1 M KOH dropwise once the sterilised medium had cooled and before pouring, until the target value was reached, the pH being monitored with pH indicator strips. Plates were inoculated with the spore suspension and incubated for 6 days; formation of a clear halo around the colony, resulting from the release of organic acids that solubilise the insoluble phosphate, was recorded as positive for phosphate solubilisation (Pikovskaya, 1948).

#### Zinc solubilisation

Zinc-solubilising ability was tested on a medium containing (g L^-1^): 10 dextrose, 1 (NH_4_)_2_SO_4_, 0.2 KCl, 0.2 K_2_HPO_4_, 0.2 MgSO_4_·7H_2_O, 1 ZnO and 15 agar. Plates were inoculated with the spore suspension and incubated for 14 days; formation of a clear halo around the colony indicated zinc solubilisation (Saravanan et al., 2004).

#### Siderophore production

it was assessed on chrome azurol S (CAS) agar following Schwyn and Neilands (1987). The assay relies on competition for iron: CAS forms a blue complex with Fe³⁺ and the detergent hexadecyltrimethylammonium bromide (HDTMA), and siderophores released by the microorganism remove iron from this complex, turning the medium orange or yellow around the colony. The HDTMA-CAS solution was prepared from three solutions: solution 1, 0.06 g CAS (Thermo Scientific, Waltham, MA, USA; cat. no. 190050250) in 50 mL distilled water; solution 2, 0.0027 g FeCl₃·6H₂O in 10 mL 10 mM HCl; and solution 3, 0.073 g HDTMA in 40 mL distilled water. Solution 1 was mixed with 9 mL of solution 2 and then with solution 3 to obtain a blue solution. The base medium contained (g L⁻¹): 4 succinic acid, 6 KH₂PO₄, 4 K₂HPO₄, 0.2 MgSO₄·7H₂O, 1 (NH₄)₂SO₄ and 20 bacteriological agar, and was adjusted to pH 6.8 with 1 M KOH. The 99 mL of HDTMA-CAS solution were then slowly added to 1000 mL of base medium cooled to approximately 50 °C until a homogeneous blue colour was obtained. Plates were inoculated with the spore suspension and incubated for 10 days; formation of an orange or yellow halo around the colony was recorded as positive for siderophore production (Schwyn and Neilands, 1987).

#### Growth on nitrogen-free medium

The ability of the fungal isolate to grow in the absence of combined nitrogen was assessed on a nitrogen-free medium following Zhang et al. (2017), containing (g L⁻¹): 10 mannitol, 5 CaCO₃, 0.3 NaCl, 0.2 KH₂PO₄, 0.2 MgSO₄·7H₂O, 0.1 CaSO₄·2H₂O and 20 agar (pH 7.2). Plates were inoculated with the spore suspension and incubated for 14 days; normal colony growth on this medium was taken as indicative of the ability to grow under nitrogen-limiting conditions.

#### Abiotic stress tolerance

Tolerance to salt stress was assessed on PDA amended with NaCl at 0, 10, 30, 60, 100, 120, 150, 300, 1000 and 3000 mM. Plates were inoculated with the spore suspension and incubated for 7 days. Colony diameter was measured in three directions (horizontal, vertical and diagonal), using the Petri dish diameter (8.5 cm) as a reference. Spores were then harvested by flooding each plate with 1 mL water and scraping the surface with an inoculation loop, and spore concentration was determined by serial dilution and haemocytometer counts (Neubauer chamber) to assess the effect of salt stress on sporulation. Three independent experiments were performed, each with three plates per concentration; colony diameter was recorded in all three experiments and conidial yield in two of them. Diameter measurements recorded as proliferated colony (a colony with non-circular growth) were treated as missing (51 of 216). To avoid pseudoreplication, the experiment mean at each concentration was taken as the unit of analysis: the mean of all valid diameter measurements (three to nine per experiment) and the mean conidial count of the three plates. For each variable, an additive linear model was fitted with NaCl concentration as a factor and experiment as a block. Colony diameter was analysed untransformed, whereas conidial yield was log₁₀-transformed to stabilise the variance, so that differences from the control are expressed as ratios. Each concentration was compared with the unamended control by a two-sided Dunnett’s test (α = 0.05), and 95% simultaneous confidence intervals were computed. Robustness was assessed with two sensitivity analyses: (i) excluding one plate in which the inoculum drop had been displaced (experiment 3, 150 mM), and (ii) using the geometric rather than the arithmetic mean of the plate counts for each experiment.

#### Substrate utilisation profiling

Carbon source utilisation was profiled on Biolog FF MicroPlates, which present 95 carbon sources and a water well in a single plate. The fungal isolate was grown on malt extract agar (MEA) plates for 10 days at laboratory room temperature. Conidia were collected from the colony surface with a sterile swab and resuspended in FF Inoculating Fluid (FF-IF; Biolog Inc.), formulated by the manufacturer for the preparation of fungal inocula. The inoculum density was adjusted spectrophotometrically to an absorbance of 0.125 at 590 nm, equivalent to the 75% transmittance recommended by the protocol, and 100 µL of the suspension was dispensed into each well. Three plates were inoculated independently, constituting the three replicates of the assay, and incubated at 26 °C in darkness for 10 days. Two readings were taken from every plate: at 750 nm, which measures turbidity and therefore mycelial growth, and at 490 nm, which measures reduction of the tetrazolium dye and therefore metabolic activity. Optical densities were recorded on a SPECTROstar Nano microplate reader (BMG LABTECH, Ortenberg, Germany). Optical densities were log-transformed to stabilise the variance and fitted by least squares as log (OD) = compound + plate, the plate term blocking the three replicates and the fit accommodating five missing readings at 750 nm. Utilisation is therefore reported as a geometric mean ratio to the water control with the plate effect removed, and values are given from the 750 nm reading unless stated otherwise. The 95 compounds were each compared with the water well by a two-sided Dunnett’s test, with the adjusted p value simulated from the observed correlation matrix of the contrasts (200,000 draws).

### Arabidopsis–fungus *in vitro* co-cultivation under salinity

Plant growth conditions, seed sterilisation and stratification, the D-root system and the co-cultivation set-up followed Gutiérrez-Manso et al. (2026), with the modifications described below. In brief, *Arabidopsis thaliana* Col-0 seeds were germinated and grown vertically *in vitro* on square Petri dishes (12 × 12 cm) containing half-strength Murashige and Skoog medium (MS; Caisson Labs; Murashige and Skoog, 1962) without vitamins or sucrose and supplemented with 300 µM KH₂PO₄ (Duchefa) and without NaCl. Co-cultivation plates contained the same medium supplemented with 100 mM NaCl. Three days after sowing, P. melinii ‘isolate 2’ was inoculated onto these plates as five 3 µL drops of a spore suspension (10³ spores mL⁻¹) placed 3 cm from the area to which seedlings would be transferred. Mock plates contained the same NaCl-supplemented medium without the fungus. Seven days after sowing, seedlings were transferred either onto the 4-day-old fungal colonies or onto mock plates, so that both inoculated and mock-inoculated plants were exposed to 100 mM NaCl only during co-cultivation. Co-cultivation lasted 10 days, after which plates were scanned. Primary root length was measured and lateral roots were counted manually on the scanned images in ImageJ (Schneider et al., 2012), total root system length was extracted from the same images with in-house software developed in our laboratory, and shoots were excised and weighed fresh. Each treatment comprised 10–19 seedlings and this experiment was performed three times independently. Measurements are expressed relative to the mean of the mock-inoculated treatment within the same experiment. Fungus-inoculated and mock-inoculated plants were compared by Student’s t-test, and differences were considered significant at P < 0.05.

## Results

In this work, we first describe the genome of *P. melinii* ‘isolate 2’ and compare it with those of other plant growth-promoting *Penicillium* species. We then examine the traits that may underlie its growth-promoting activity, combining carbon source profiling, *in vitro* assays and genome mining, and finally ask whether the isolate tolerates salt and promotes plant growth under salinity.

### Genome assembly and nuclear genome architecture

Four assembly strategies were compared, three from ONT long reads alone (Flye, Raven and Miniasm) and one hybrid assembly of ONT and Illumina reads (MaSuRCA), all in the range 29.2–29.5 Mb, consistent with the previously reported genome size for *P. melinii* ‘isolate 2’ (Gutiérrez-Manso et al., 2026). MaSuRCA yielded the most contiguous initial assembly (19 contigs) and was selected for downstream scaffolding. Two sequential rounds of reference-guided scaffolding with RagTag, followed by long-read scaffolding with ntLink, progressively consolidated the 19 contigs into 9 scaffolds, with a mean GC content of 50%. Identification and removal of three mitochondrial scaffolds, distinguished by their markedly lower mean GC content (25.81%) relative to the nuclear scaffolds, yielded a final nuclear assembly of six scaffolds (Table 1; Figure 1A). BlobTools analysis revealed no evidence of contamination: the assembly formed a single cluster of uniform GC content (∼0.5) and coverage, with 50.3% of mapped sequence assigned to *Penicillium atrosanguineum* and all other taxa below 2%, consistent with a single-organism assembly. BUSCO completeness (ascomycota_odb12, n = 2,826) was assessed at each processing step. The initial MaSuRCA assembly yielded 97.5% complete BUSCOs (single-copy: 96.5%; duplicated: 1.0%; fragmented: 0.3%; missing: 2.2%). Scores remained stable across Illumina polishing, RagTag, and ntLink scaffolding steps (97.6% complete; fragmented: 0.3%; missing: 2.1%). The final six-scaffold nuclear assembly achieved 97.6% complete BUSCOs (single-copy: 96.6%; duplicated: 1.0%; fragmented: 0.3%; missing: 2.1%), confirming that no gene content was lost at any processing step (Figure 1B; Table 1).

**Table 1.** Summary statistics for the *Penicillium melinii* ‘isolate 2’ nuclear genome assembly and structural annotation. BUSCO run with ascomycota_odb12 (n = 2,826 BUSCOs).

| **Assembly** |  |
|---|---|
| Total length | 29.39 Mb |
| Scaffolds | 6 |
| N50 | 9.29 Mb |
| L50 | 2 |
| Largest scaffold | 10.94 Mb |
| GC content | 50.0 % |
| N's per 100 kbp | 0.95 |
| BUSCO completeness (ascomycota_odb12) | 97.6 % |
| **Structural annotation** |  |
| Protein-coding genes | 9,566 |
| tRNA genes | 260 |
| Complete CDS (start + stop) | 9,550 (99.8 %) |
| Mean gene length | 1,620 bp |
| Mean protein length | 491 aa |
| Total exons | 28,654 |
| Mean exons per mRNA | 3.0 |
| Mean exon length | 492 bp |
| Mean intron length | 72 bp |
| Single-exon genes | 1,851 (19.3 %) |

**Figure 1.**
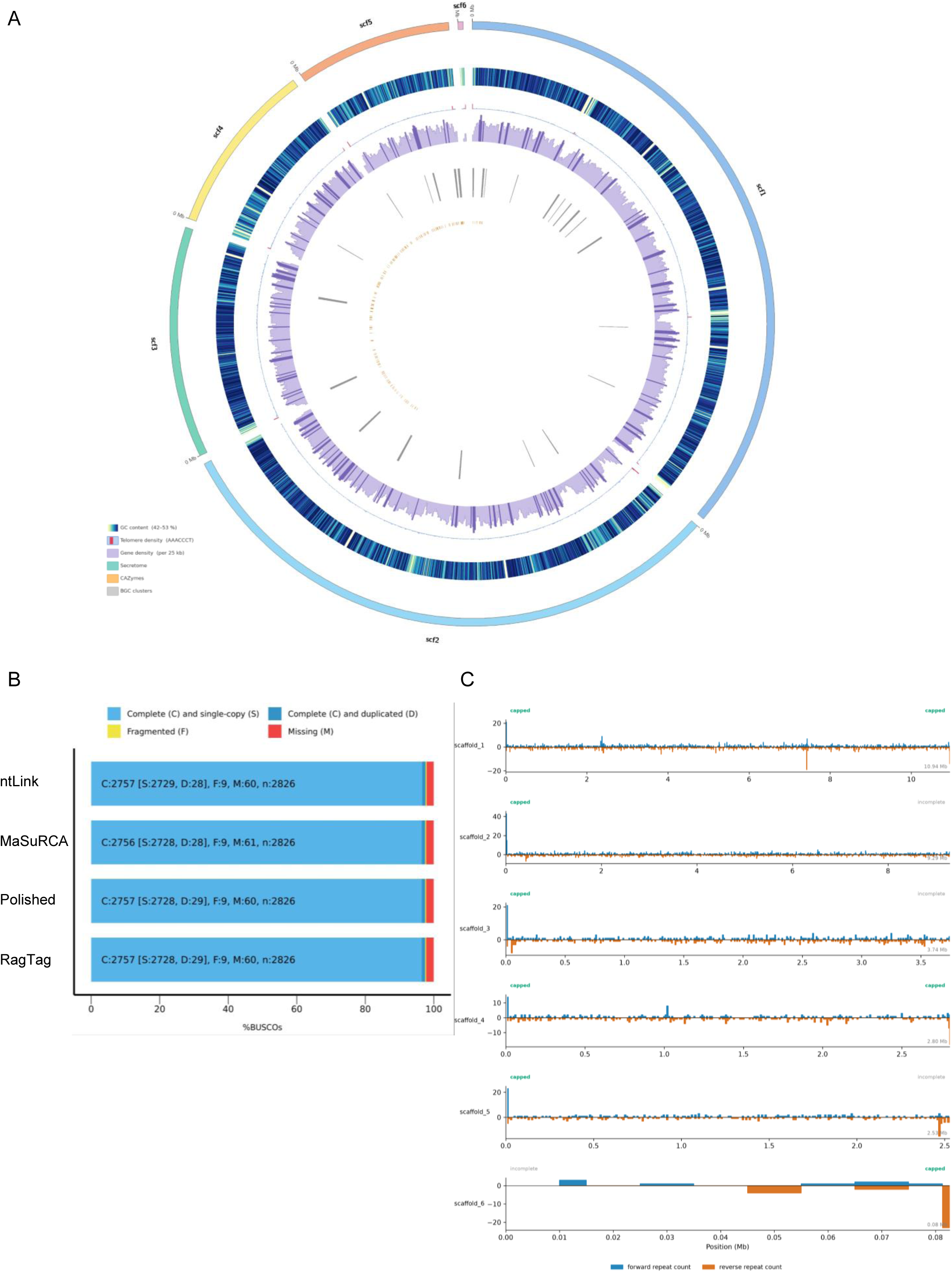
Nuclear genome assembly of *P. melinii* ‘isolate 2. (A) Circos plot of the nuclear assembly (six scaffolds, 29.39 Mb). Tracks from outermost to innermost: (1) scaffold ideogram (scf1–scf6), coloured by scaffold; (2) GC content (42–53%, YlGnBu gradient, 10 kb windows); (3) Telomere repeat frequency across the six nuclear scaffolds of Penicillium melinii isolate 2. The repeat motif AAACCCT (4,117 occurrences); (4) gene density (per 25 kb window, purple histogram); (5) secretome proteins (teal ticks); (6) CAZymes (orange ticks); (7) biosynthetic gene cluster (BGC) regions (grey bars). (B) BUSCO completeness (ascomycota_odb12, n = 2,826) of the assembly at successive processing steps. (C) Telomere repeat frequency across the six nuclear scaffolds, identified with tidk; peaks at scaffold termini indicate telomere-associated sequence.

The predominant telomeric repeat unit was identified as AAACCCT/ TTTAGGG (4,117 occurrences), a 7-mer distinct from the TTAGGG motif typical of filamentous ascomycetes. Telomeric signal was detected at both termini of scaffold_1 and scaffold_4 and at one terminus of the remaining four scaffolds; internal peaks at scaffold_1 (∼7.4 Mb) and scaffold_4 (∼1.0 Mb) coincide with inter-scaffold junctions, likely marking former contig ends. Overall, two of six scaffolds (33%) are fully telomere-capped, supporting a near-chromosome-level assembly (Figure 1C).

Structural annotation predicted 9,826 gene models on the *P. melinii* ‘isolate 2’ nuclear genome, comprising 9,566 protein-coding loci and 260 tRNA genes, with no alternative transcripts detected. Of the coding models, 9,550 (99.8%) were complete with both start and stop codons. The combined use of the species-specific *Penicillium* NR protein database and cross-species RNA-seq alignments (9,566 models) achieved 98.7% complete BUSCOs (single-copy: 97.9%; fragmented: 0.5%; missing: 0.8%), a 2.9 percentage point improvement in completeness and an approximately four-fold reduction in fragmented gene models relative to the other strategies, and was therefore selected as the final annotation (Figure 2A).

**Figure 2.**
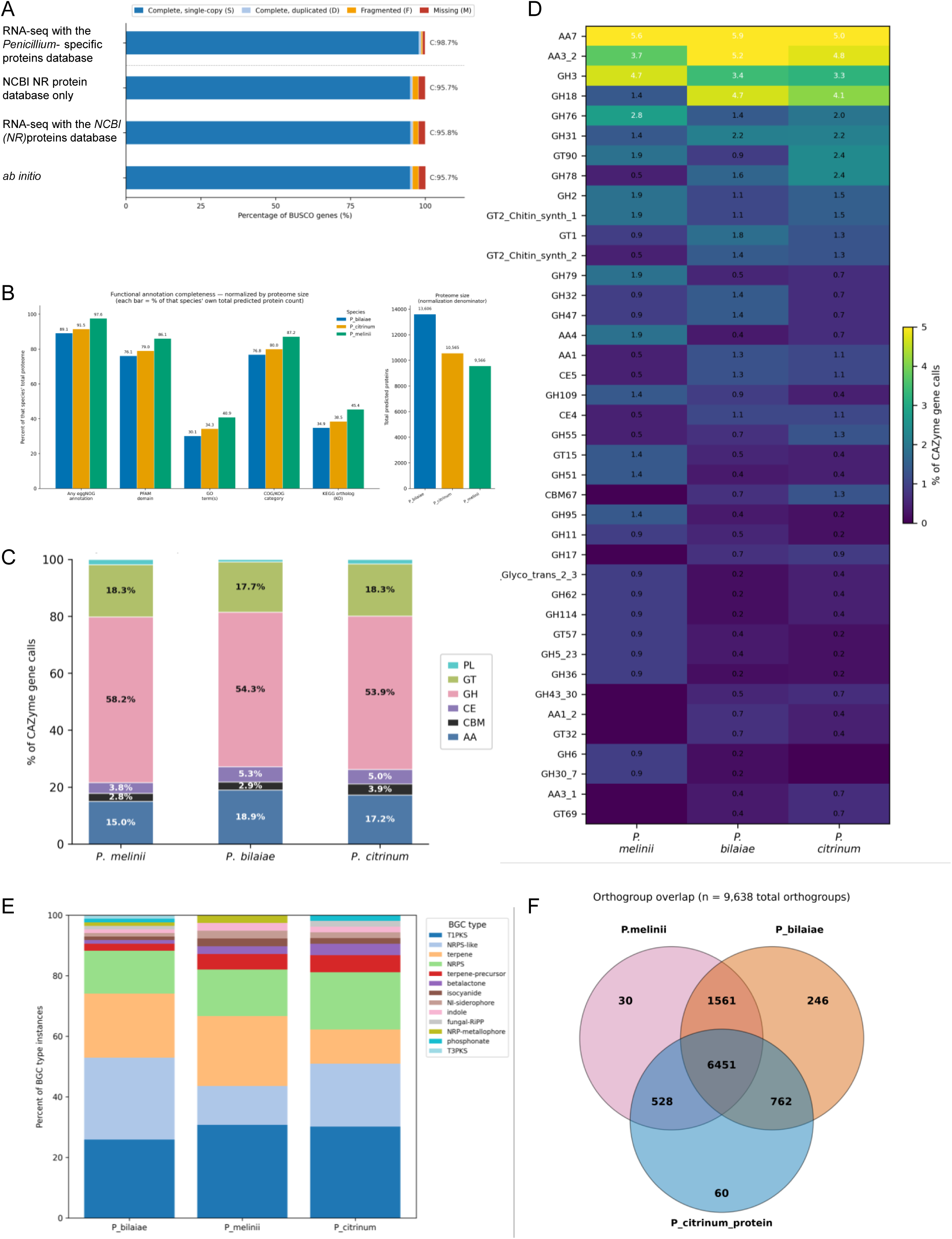
Annotation quality, carbohydrate-active enzyme repertoire and genomic context of *Penicillium melinii* ‘isolate 2’ alongside two congeners, *P. bilaiae* ATCC 20851 and *P. citrinum* B9. (A) BUSCO completeness (ascomycota_odb12, n = 2,826) of the gene models of *P. melinii* ‘isolate 2’ obtained with four annotation strategies: no external evidence (ab initio), RNA-seq evidence and non redundant (NR) proteins database from NCBI, NCBI (NR) protein evidence only, and *Penicillium* species-specific protein database plus RNA-seq evidence (selected); complete single-copy (dark blue), complete duplicated (light blue), fragmented (yellow) and missing (red). (B) Functional annotation completeness, each bar the percentage of that species’ own predicted proteins carrying an annotation of the given type; the inset gives the proteome used as denominator (*P. bilaiae*, 13,606; *P. citrinum*, 10,565; *P. melinii*, 9,566). *P. melinii* has the highest coverage in all five categories while having the smallest proteome. (C) CAZyme class composition (dbCAN v10; *P. melinii*, n = 207; *P. bilaiae*, n = 535; *P. citrinum*, n = 443). Glycoside hydrolases dominate in all three (52.8–57.5%), and *P. melinii* has the smallest share of auxiliary activities (15.5%). (D) The 40 CAZyme families of highest cross-species variance; cell values and colour give each family’s share of that species’ CAZyme gene calls. Dark cells mark families with no dbCAN call, among them CBM67, GH17, GH43_30, AA1_2 (ferroxidases; a FET3 homologue, FUN_001010, is present but received no CAZyme assignment) and AA3_1 (cellobiose dehydrogenases) in *P. melinii*. (E) Biosynthetic gene cluster composition (antiSMASH v8.0, relaxed strictness), each bar the proportional distribution of cluster types within that species’ own complement. (F) Orthogroup overlap inferred by OrthoFinder v3.1.2 from the same predicted proteomes; numbers are orthogroup counts per intersection and sum to the 9,638 total. Circles are drawn at equal size and are not proportional to those counts. *P. melinii* has the fewest orthogroups of its own, 30 against 246 and 60.

Functional annotation supported the quality of the gene prediction: 97.6% of predicted proteins (n = 9,340) received at least one annotation by eggNOG-mapper, of which 86.1% (n = 8,233) carried at least one Pfam domain assignment. Of all predicted proteins, 2.2% (n = 207) were annotated as CAZymes and 3.4% (n = 325) as proteases.

### Mitochondrial genome assembly and annotation

High-molecular-weight DNA sequencing and multi-strategy assembly together enabled recovery of complete mitochondrial contigs alongside the nuclear genome; candidate mitochondrial sequences were distinguished by their elevated read coverage and reduced GC content relative to nuclear contigs and confirmed by homology-based classification. The *P. melinii* ‘isolate 2’ mitochondrial genome assembled as a single circular contig of 26,730 bp (Table 2; Supplementary Figure 1), comparable with published *Penicillium* sp. mitogenomes (25.5–28.6 kb) (Jiang et al., 2020; Malik et al., 2024). Circularity was confirmed by uniform read coverage across the genome (mean 3,190×; terminal 20-bp windows 3,126× and 3,124×, with no coverage drop at the sequence termini) and by 3,187 of 29,232 alignments spanning the sequence join point without interruption, directly confirming physical continuity of the two ends.

**Table 2.** Summary statistics for the *Penicillium melinii* ‘isolate 2’ mitochondrial genome assembly and annotation.

| **Assembly** |  |
|---|---|
| Total length | 26,730 bp |
| Topology | Circular |
| **MITOS2 annotation** |  |
| Total annotated features | 39 |
| Protein-coding genes | 15 |
| rRNA subunits | 2 (rrnS 1,450 bp; rrnL 2,258 bp) |
| tRNA genes | 22 (all 20 standard amino acids) |
| Coding density | 72.2 % (19,300 / 26,730 bp) |
| Intron-containing genes | 0 |
| Strand orientation | Single strand |

Annotation with MITOS2 identified 39 genetic elements: 15 protein-coding genes (*atp6*, *atp8*, *atp9*, *cob*, *cox1*, *cox2*, *cox3*, *nad1*, *nad2*, *nad3*, *nad4*, *nad4L*, *nad5*, *nad6*, *rps3*), two ribosomal RNA genes (*rrnS*, 1,450 bp; *rrnL*, 2,258 bp), and 22 transfer RNA genes covering all 20 standard amino acids, with duplicated isoacceptors for leucine (*trnL1*, *trnL2*) and serine (*trnS1*, *trnS2*), as is typical in fungal mitogenomes. All genes were encoded on the same strand and no intron-containing gene structures were detected. Coding sequences occupied 19,300 of 26,730 bp (72.2% coding density), consistent with the compact organisation and limited intergenic regions typical of fungal mitogenomes (Supplementary Figure 1). The *cox1* gene (1,611 bp; coordinates 25,696–27,306) spans the linearisation point of the circular assembly and was annotated as a single continuous open reading frame in the final circular map. Complete mitochondrial genomes remain limited across the *Penicillium* genus (Joardar et al., 2012; Malik et al., 2024), and this assembly represents, to the best of our knowledge, the first complete mitogenome reported for *P. melinii*, expanding the genomic resources available for comparative studies of plant growth-promoting *Penicillium* species.

### Phylogenomic placement of *P. melinii* ‘isolate 2’

To place this genome in context, we first confirmed the phylogenetic position of ‘isolate 2’ within the genus. The identity of the ‘isolate 2’ was assigned by Gutiérrez-Manso et al. (2026) from a multi-locus analysis; the complete nuclear genome assembly reported here allows that assignment to be tested at genome scale. A phylogenomic supermatrix was constructed by extracting single-copy BUSCO orthologs (eurotiales_odb10) present in *P. melinii* ‘isolate 2’ and 16 reference taxa (15 Penicillium spp. and Aspergillus fumigatus as outgroup) at 100% occupancy (BUSCO_phylogenomics; Supplementary Figure 2). Individual gene trees were estimated with FastTree and the concatenated supermatrix alignment was used to infer the final species topology under approximate maximum likelihood (FastTree; SH-like local support values). *Penicillium melinii* ‘isolate 2’ was placed in a fully supported clade (SH = 1.0) with *P. maclennaniae* as its immediate sister taxon, and *P. atrosanguineum* and *P. corylophilum* as successive relatives. *Penicillium citrinum* B9 resolved within the same broader clade, indicating moderate phylogenetic proximity to *P. melinii* ‘isolate 2’ (Supplementary Figure 2). In contrast, *P. bilaiae* ATCC 20851 was placed sister to the clade comprising *P. citrinum*, *P. janthinellum*, *P. oxalicum*, and the *P. melinii* lineage, confirming deep divergence from *P. melinii* ‘isolate 2’ despite their shared PGP phenotype (Supplementary Figure 2). *Penicillium rubens*, *P. griseofulvum*, *P. olsonii*, and *P. canescens* resolved in a distal clade alongside *P. expansum*, *P. italicum*, *P. digitatum*, and *P. chrysogenum*, phylogenetically distant from the *P. melinii* lineage. This topology agrees with the multi-locus assignment of Gutiérrez-Manso et al. (2026) and confirms it at genome scale.

### Comparative genomics of plant growth-promoting Penicillium species

To establish whether the predicted proteome of *P. melinii* ‘isolate 2’ (9,566 proteins) is distinctive among PGP *Penicillium* species or representative of the genus, we applied the same functional annotation pipeline to two species already reported to benefit plant growth, *P. citrinum* B9 (10,565 proteins) and *P. bilaiae* ATCC 20851 (13,606). Annotation coverage was high in all three proteomes and, despite having the smallest proteome, *P. melinii* ‘isolate 2’ returned the highest rates in all five categories (*P. melinii* vs. *P. citrinum* and *P. bilaiae*): eggNOG-mapper hits (97.6% vs. 91.5% and 89.1%), Pfam domains (86.1% vs. 79.0% and 76.1%), COG/KOG categories (87.2% vs. 80.0% and 76.8%), KEGG orthologs (45.4% vs. 38.5% and 34.9%) and GO terms (40.9% vs. 34.3% and 30.1%; Figure 2B). Supplementary Tables 1-3 list only the proteins with an eggNOG-mapper hit (9,340, 9,672 and 12,127, respectively).

To focus the comparative analysis on the functional repertoire underlying plant growth promotion, we next examined secreted protein candidates across all three *Penicillium* species, each previously reported as beneficial to plant growth. *Penicillium melinii* ‘isolate 2’ harboured a contracted effector repertoire relative to its secretome size compared with both reference species. The consensus secretome comprised 548 proteins in *P. melinii* ‘isolate 2’, 805 in *P. bilaiae* ATCC 20851, and 516 in *P. citrinum* B9, of which 471 (85.9%), 718 (89.2%) and 459 (88.9%) were predicted to localise extracellularly by DeepLoc v0.1, respectively (Supplementary Tables 1–3). EffectorP3 classified 112 *P. melinii* ‘isolate 2’ proteins as candidate effectors, 83 apoplastic and 11 cytoplasmic, the smallest complement among the three species despite an extracellular protein count comparable to P. citrinum B9 (129 candidate effectors) and markedly below that of P. bilaiae ATCC 20851 (185) (Supplementary Tables 1–3). The smaller effector complement is not offset by a larger CAZyme or BGC repertoire (see below), which points to an overall reduction of the secreted repertoire rather than to a shift between protein classes.

A total of 207, 535, and 443 CAZyme-encoding genes were identified in *P. melinii* ‘isolate 2’, *P. bilaiae* ATCC 20851 and *P. citrinum* B9, representing 2.2%, 3.9%, and 4.2% of each predicted proteome, respectively (Figure 2C). *Penicillium melinii* thus harbours the smallest CAZyme repertoire both in absolute and relative terms, which may reflect its more specialised ecological niche compared to the broader saprotrophic lifestyles of the other two species. Despite differences in CAZymes repertoire size, the three species showed highly conserved class-level composition (Figure 2C). Glycoside hydrolases (GH) were the dominant class in all three species (58.2%, 54.3%, and 53.9% of class assigments per gene, respectively. Glycosyltransferases (GT; ∼18% across all species) and Auxiliary Activities (AA; 15.0–18.9%) were the next most abundant classes. Carbohydrate esterases (CE; 3.8–5.3%), Carbohydrate-Binding Modules (CBM; 2.8–3.9%), and Polysaccharide Lyases (PL; 0.9–1.9%) were consistently minor components (Figure 2C).

At the family level, the CAZyme families with the highest representation across the three species included AA7, AA3_2, GH3, GH18 and GH76 (Figure 2D). The most marked difference was the contraction of GH18 chitinases in *P. melinii* ‘isolate 2’ (1.4% of CAZyme gene calls, against 4.7% in *P. bilaiae* and 4.1% in *P. citrinum*). AA3_2 (aryl-alcohol oxidases and glucose oxidases) was most represented in *P. bilaiae* (5.2%) and least in *P. melinii* (3.7%). *P. melinii* ‘isolate 2’ lacked detectable representatives of AA3_1 (cellobiose dehydrogenases), GH17 (β-1,3-glucanases), CBM67 and GH43_30, which were present in both congeners. No gene was assigned to AA1_2 (ferroxidases), although the genome encodes a FET3 homologue (FUN_001010) that received no CAZyme assignment. In *P. melinii* ‘isolate 2’, 39.6% (82 of 207) of CAZyme-encoding genes are predicted to be secreted comparable to *P. bilaiae* (42.2%) and exceeding *P. citrinum* (29.1%).

Biosynthetic gene clusters (BGCs) encode the secondary metabolites that mediate nutrient mobilisation (Haas, 2014), pathogen suppression (Keller et al., 2005), and direct phytostimulation at the plant–fungal interface (Vinale et al., 2012). Comparative genome mining was therefore used to delineate the conserved, shared, and species-specific components of the biosynthetic repertoire across the three PGP *Penicillium* proteomes. Although *P. melinii* ‘isolate 2’ harboured the fewest BGCs among the three species, the distribution of cluster types was broadly similar across all three proteomes, suggesting a quantitative contraction rather than qualitative divergence or the emergence of species-specific biosynthetic pathways. Genome mining with antiSMASH predicted 37 BGC regions in *P. melinii* ‘isolate 2’, compared with 78 in *P. bilaiae* ATCC 20851 and 44 in *P. citrinum* B9 (Figure 2E). The most frequent BGC types were T1PKS (n = 10), terpene (n = 10, including two terpene-precursor clusters), and NRPS or NRPS-like (n = 8); four hybrid clusters and five additional types (betalactone, isocyanide, NI-siderophore, indole, and NRP-metallophore) completed the predicted repertoire. A comparison with the MIBiG reference database identified 12 regions with known compound similarity: five at high confidence, comprising two independent terpene clusters matching clavaric acid (regions 1.5 and 1.10), a terpene–T1PKS hybrid cluster for andrastin A, a T1PKS cluster for fusarin C, and an NRPS-like cluster for choline; three at medium confidence (HEx-tc1 terpene, nidulanin A, and squalestatin S1); and four at low confidence (aurofusarin, bikaverin, harziphilone, and paraherquamide).

To identify conserved and species-specific components of the plant growth-promoting repertoire among the three species, we compared the three predicted proteomes through orthologous group inference using OrthoFinder. The analysis identified 9,638 orthogroups (Figure 2F), with 89.3% of *P. melinii* ‘isolate 2’ predicted proteins assigned to at least one orthogroup (mean orthogroup size 3.1 genes). Among the 9,638 orthogroups, 6,451 were shared by all three species, representing the conserved core of this *Penicillium* lineage (Figure 2F). An additional 1,561 orthogroups were shared exclusively between *P. melinii* ‘isolate 2’ and *P. bilaiae* ATCC 20851, 528 between *P. melinii* ‘isolate 2’ and *P. citrinum* B9, and 762 between the two reference species. *P. melinii* ‘isolate 2’ harboured 30 species-specific orthogroups containing 70 genes, markedly fewer than *P. bilaiae* ATCC 20851 (246 orthogroups) or *P. citrinum* B9 (60 orthogroups). Among these 70 species-specific proteins (Supplementary Table 4), 59 (84.3%) received functional annotations via eggNOG-mapper; 8 were predicted to localise extracellularly by DeepLoc v0.1, and one (FUN_005453; 72 aa) was identified as a candidate effector by EffectorP3. Sequence similarity searches against the NCBI NR database (blastp) returned hits exclusively in *Penicillium* species (best hit: *P. atrosanguineum*, 100% identity, E = 6 × 10⁻⁴¹), with a distant match to a glycoside hydrolase family 130 (GH130) protein; no conserved functional domain could be assigned.

### Traits underlying plant growth promotion and their genomic candidates

‘Isolate 2’ promotes the growth of several plant species (Gutiérrez-Manso et al., 2026), but the fungal traits behind this activity are unknown. We approached them at three levels: the carbon sources the isolate can use, the plant growth-promoting and hydrolytic activities it displays *in vitro*, and the genes that could account for them.

First, to define the nutritional strategy of ‘isolate 2’, the phenotypic counterpart of the CAZyme repertoire described above, carbon source utilisation was profiled on Biolog FF MicroPlates as previously applied to plant-associated fungi (Hochenegger et al., 2025). Utilisation of each compound is expressed as fold change, the optical density at 750 nm relative to that of the water well of the same plate, so that a value of 1 is no growth above the control. *P. melinii* ‘isolate 2’ utilised all 95 carbon sources tested, with 63 significantly above the water control (median 2.88-fold; 28 compounds ≥ 4-fold; 18 > 6-fold; Figure 3, Supplementary Table 5). Carbohydrates dominated the substrate ranking: oligosaccharides and polysaccharides ranked first (median 5.19), monosaccharides second (5.07), followed by polyols (3.06), aryl and methyl glycosides (2.88), amino acids and peptides (2.50), amino sugars (2.02) and organic acids (1.91). The six highest individual values were the disaccharides and oligosaccharides maltose (8.98), palatinose (8.95), cellobiose (8.64), β-methyl-D-glucoside (8.41), gentiobiose (8.31) and maltotriose (8.11), with the monosaccharides α-D-glucose (7.83), mannose and ribose (7.46 each) and xylose (6.96) among the next-ranked compounds. Twelve compounds produced a metabolic signal without concomitant growth; ten were organic acids or amino acid derivatives, including fumaric, succinic, α-ketoglutaric, L-malic and β-hydroxybutyric acids (Supplementary Table 5).

**Figure 3.**
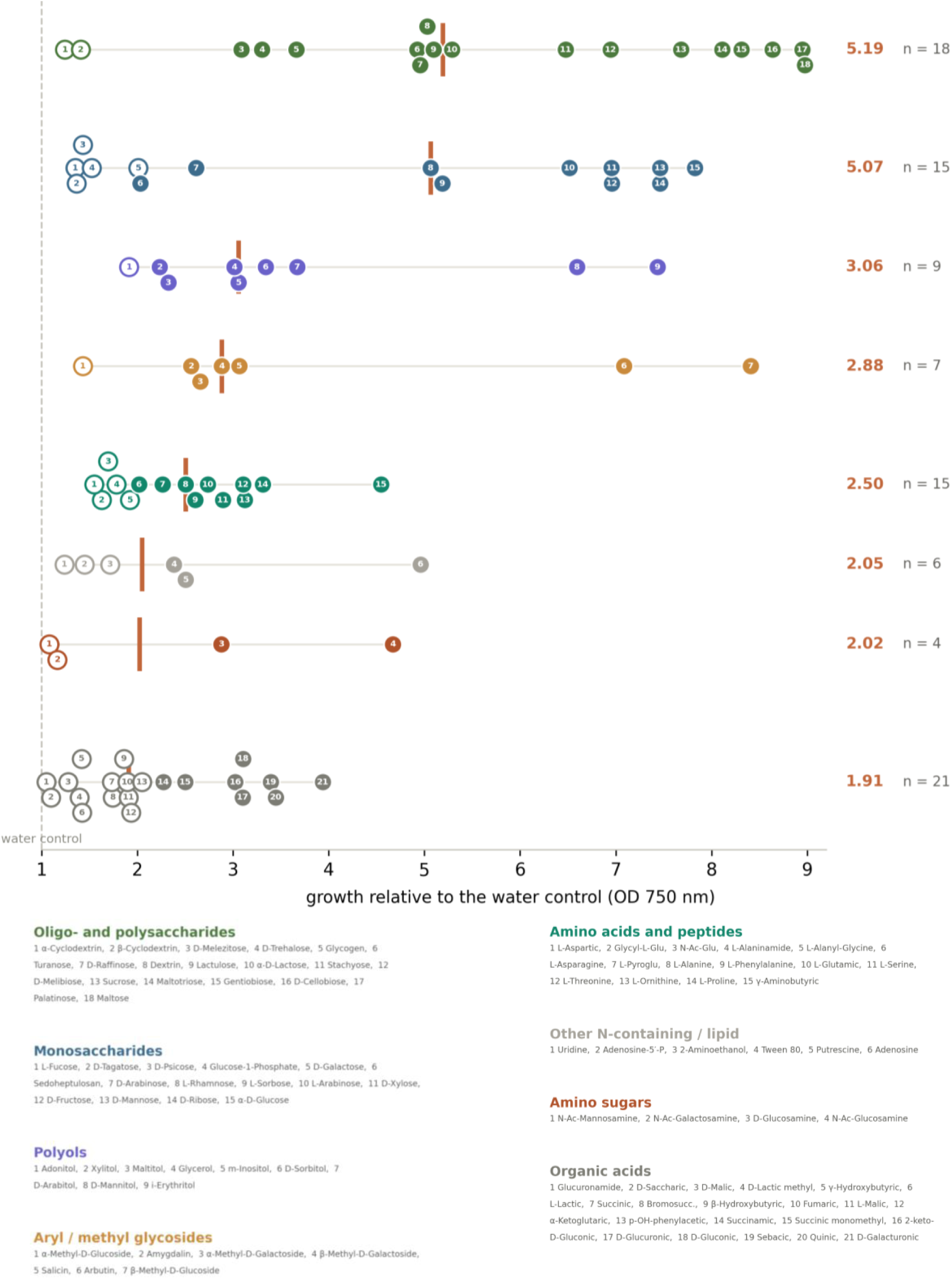
Substrate utilisation of *Penicillium melinii* ‘isolate 2’ across the 95 carbon sources of the Biolog FF MicroPlate, grouped by substrate class. Each numbered marker is one compound; within each class, numbering runs from the lowest to the highest value and the key is given beneath the class name. Growth is the optical density at 750 nm expressed relative to the water control (dashed line at 1), estimated from three independently inoculated plates as a geometric mean ratio with the plate effect removed. Filled markers differ significantly from the water control (two-sided Dunnett’s test, p < 0.05) and open markers do not; with three plates the design resolves ratios of about 2 and above, so an open marker means unresolved rather than unused. The vertical bar marks the class median, given on the right with the number of compounds in the class. Compound-level values, adjusted p values and the corresponding readings at 490 nm are given in Supplementary Table 5.

To gain insight into the mechanisms by which *P. melinii* ‘isolate 2’ may exert its beneficial effect on plant growth, the isolate was screened *in vitro* for a panel of traits associated with plant growth promotion. The isolate was positive for 7 of the 11 traits assayed (Figure 4). Because a single isolate was characterised, all assays were scored qualitatively as presence or absence of the diagnostic colour change or halo; halo diameters and colorimetric values were not quantified, since such measurements are informative when comparing several isolates. Most of the positive traits were related to nutrient acquisition and mobilisation. The isolate produced siderophores on chrome azurol S (CAS) agar: by 10 days post-inoculation (dpi), the blue CAS–Fe³⁺–HDTMA complex was decolourised around the colonies, leaving a yellow-orange halo (Figure 4A). On Pikovskaya (PVK) agar containing tricalcium phosphate, clear halos developed around the colonies on media adjusted to starting pH values of 5.7, 7.0 and 8.0 (Figure 4B); because the medium is unbuffered, this does not by itself show solubilisation at alkaline pH (see Discussion). The isolate also solubilised zinc, as shown by the clearing halo that surrounded the colonies on ZnO-amended agar at 14 dpi (Figure 4C). Ammonia production was detected in peptone water at 3 dpi, when Nessler’s reagent turned the cultures orange-brown whereas uninoculated controls remained yellow (Figure 4D). Urease activity was time-dependent: on urea agar with phenol red, the medium remained yellow at 7 dpi but had turned pink-fuchsia by 14 dpi, consistent with urease activity, although ammonia released from the peptone in the medium may also contribute to the alkalinisation (Figure 4E). In contrast, the isolate did not grow under nitrogen limitation; on nitrogen-free medium mannitol medium, growth was restricted to the inoculation spotat 14 dpi, and the isolate was therefore scored negative for nitrogen oligotrophy (Figure 4F).

**Figure 4.**
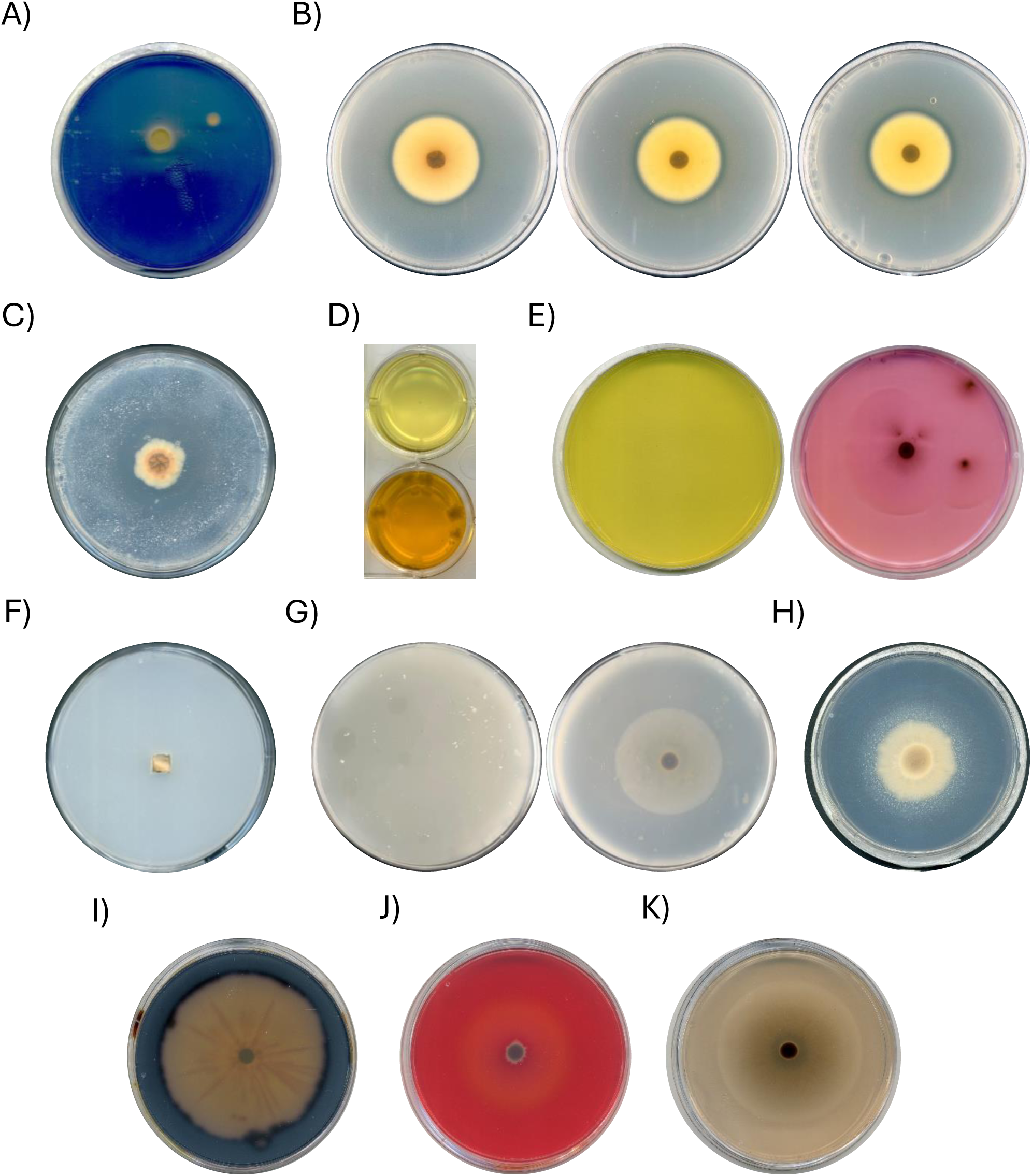
*In vitro* plant growth-promoting traits of *Penicillium melinii* ‘isolate 2’. (A) Siderophore production on chrome azurol S (CAS) agar at 10 dpi; the blue medium turns yellow-orange around the colony. (B) Solubilisation of tricalcium phosphate on Pikovskaya agar adjusted to starting pH values of 5.7, 7.0 and 8.0 (left to right); clear halos surround the colonies. (C) Solubilisation of zinc oxide at 14 dpi, shown by the clear halo around the colony. (D) Ammonia production in peptone water at 3 dpi after addition of Nessler’s reagent: uninoculated control (top, yellow) and inoculated culture (bottom, orange-brown). (E) Urease test on urea agar with phenol red at 7 dpi (left, yellow) and 14 dpi (right, pink-fuchsia). (F) Nitrogen-limited mannitol medium at 14 dpi; growth is restricted to the inoculation spot. (G) Gelatine hydrolysis at 14 dpi after flooding with saturated ammonium sulphate: uninoculated control (left) and inoculated plate (right), with a clear zone around the colony against the turbid background of precipitated gelatine. (H) Lipolytic activity on Tween 20 agar with CaCl₂ at 14 dpi, shown by the opaque halo of precipitated calcium salts around the colony. (I) Starch agar stained with iodine at 14 dpi; the medium stains dark blue-black with no clear zone around the colony. (J) Carboxymethylcellulose agar stained with Congo red at 10 dpi; no clearing zone forms around the colony. (K) 1-Naphthol agar at 14 dpi; no blue oxidation product forms.

The isolate also displayed extracellular hydrolytic activities, although these were restricted to proteins and lipids. Proteolytic activity was detected on gelatine agar: after flooding with saturated ammonium sulphate, a clear zone surrounded the colony against a turbid background of precipitated gelatine at 14 dpi (Figure 4G). Lipolytic activity was detected on Tween 20 agar supplemented with CaCl₂; at 14 dpi, an opaque halo of precipitated calcium salts of the released fatty acids surrounded the colonies, whereas the uninoculated medium remained transparent (Figure 4H). No amylase, cellulase or laccase activity was detected: on GYP agar with soluble starch, iodine (I₂/KI) stained the medium uniformly dark blue-black with no clear zone around the colony at 14 dpi (Figure 4I); on carboxymethylcellulose (CMC) agar stained with Congo red, no clearing was observed around the colony at 10 dpi (Figure 4J); and on 1-naphthol agar, no blue oxidation product formed at 14 dpi (Figure 4K).

Overall, the *in vitro* PGP profile of *P. melinii* ‘isolate 2’ is dominated by nutrient-mobilising traits (siderophore production, phosphate and zinc solubilisation, ammonia release and urease activity) together with proteolytic and lipolytic activities, whereas the isolate lacked detectable amylase, cellulase and laccase activities and was unable to grow under nitrogen limitation.

To link these activities to the genome, we searched the functional annotation for candidate genes, both for the traits detected in the plate assays and for mechanisms of plant growth promotion that these assays do not capture (Table 3). Candidates were found for most positive assays of nutrient mobilisation. For siderophore production (Figure 4A), the L-ornithine N⁵-monooxygenase SidA (FUN_005658) lies next to a siderophore NRPS (FUN_005657) within the same predicted BGC region, and a second locus contains three contiguous genes similar to *sidH* (FUN_006196), *sidF* (FUN_006197) and *sidD* (FUN_006199, an NRPS fragment); three siderophore–iron transporters (Sit1, FUN_005561; FUN_001365; MirB, FUN_005646) and the reductive uptake pair Ftr1–Fet3 (FUN_001009, FUN_001010) are also encoded. For tricalcium phosphate solubilisation (Figure 4B), the candidates are genes of organic acid metabolism: three citrate synthases (CIT1, FUN_000550; CIT3, FUN_000965; FUN_005306), two isocitrate lyases (ICL1, FUN_003380; ICL2, FUN_000937) and malate synthase (FUN_000949). Although the isolate solubilised ZnO *in vitro* (Figure 4C), no secreted protein specifically associated with zinc solubilisation was identified, and the zinc-related genes found encode ZIP- and Nramp-family importers involved in uptake rather than mobilisation; ZnO solubilisation may therefore rely on the same organic acid secretion proposed for phosphate. Eight phosphatases with predicted signal peptides could act on organic phosphorus, which was not assayed: three histidine acid phosphatases including PHO5 (FUN_006111, FUN_001989, FUN_006396), three PhoA-type acid phosphatases (FUN_008186, FUN_009045, FUN_005393), a purple acid phosphatase (FUN_000884) and an alkaline phosphatase (FUN_004427). Urease activity (Figure 4E) is matched by URE1 (FUN_006517), and gelatine hydrolysis (Figure 4G) and ammonia release from peptone (Figure 4D) by 16 secreted proteases, among them a PRB1-like subtilisin (FUN_000372), the aspartic proteases PepA (FUN_003513, FUN_005935) and an M35 metalloprotease (FUN_009712).

**Table 3.** Candidate genes for salt tolerance and plant growth-promoting traits in the *Penicillium melinii* ‘isolate 2’ genome, with the phenotype of the isolate to which each relates. Gene IDs are funannotate gene models; SP, predicted signal peptide; Eff, predicted effector (EffectorP3); NRPS, nonribosomal peptide synthetase; PLP, pyridoxal phosphate.

| Trait | Gene product | Gene ID |
|---|---|---|
| **Iron** | L-ornithine N<sup>5</sup>-monooxygenase SidA; adjacent siderophore NRPS | FUN_005658; FUN_005657 |
|  | SidH-, SidF- and SidD-like, contiguous (SidD-like is an NRPS fragment) | FUN_006196, FUN_006197, FUN_006199 |
|  | Siderophore–iron transporters (Sit1, MirB); reductive uptake (Ftr1, Fet3) | FUN_005561, FUN_001365, FUN_005646; FUN_001009, FUN_001010 |
| **Phosphorus (mineral), zinc** | Citrate synthases (CIT1, CIT3); isocitrate lyases (ICL1, ICL2); malate synthase | FUN_000550, FUN_000965, FUN_005306; FUN_003380, FUN_000937; FUN_000949 |
| **Phosphorus (organic)** | Histidine acid phosphatases incl. PHO5; PhoA-type acid phosphatases; purple acid phosphatase; alkaline phosphatase (all SP) | FUN_006111, FUN_001989, FUN_006396; FUN_008186, FUN_009045, FUN_005393; FUN_000884; FUN_004427 |
| **Nitrogen** | Urease URE1 | FUN_006517 |
|  | Secreted proteases: subtilisin PRB1, aspartic PepA, M35 metalloprotease (SP) | FUN_000372; FUN_003513, FUN_005935; FUN_009712 |
| **Ethylene** | ACC deaminase (EC 3.5.99.7); PLP-dependent enzyme (ACC deaminase or D-cysteine desulfhydrase) | FUN_006661; FUN_000448 |
| **Auxin** | Tryptophan biosynthesis (TRP2, TRP4, TRP3, tryptophan synthase) | FUN_003446, FUN_001950, FUN_004234, FUN_003233, FUN_003573 |
|  | Indole-3-pyruvate route: aromatic aminotransferase ARO8 | FUN_003207 |
|  | Tryptamine route: aromatic L-amino acid decarboxylase | FUN_009706 |
|  | Indole-3-acetonitrile route: generic nitrilases | FUN_003565, FUN_006226 |
| **Cytokinin** | tRNA-isopentenyltransferase | FUN_004731 |
|  | LOG-like phosphoribohydrolases | FUN_003275, FUN_004765 |
| **Colonisation** | LysM proteins (SP; FUN_008586 and FUN_009355 also Eff) | FUN_001690, FUN_008586, FUN_009355 |
|  | Class I hydrophobins (SP except FUN_004030); class II hydrophobins (SP) | FUN_000077, FUN_000268, FUN_000429, FUN_004030; FUN_009018, FUN_009291 |
|  | Secreted GH10 xylanase; secreted AA9 lytic polysaccharide monooxygenases (SP) | FUN_009088; FUN_005814, FUN_009388 |
| **Induced resistance** | Cerato-platanin (SP, Eff) | FUN_002426 |
| **Na<sup>+</sup> efflux** | ENA-type P-type ATPases (two ENA2, one ENA-type, one Cta3-like) | FUN_004879, FUN_009227, FUN_007340, FUN_002495 |
| **Na<sup>+</sup>/H<sup>+</sup> antiport** | Nha1-type (plasma membrane); NHX1-type (endosomal) | FUN_005033; FUN_000670 |
| **Vacuolar cation sequestration** | Cation/H<sup>+</sup> exchanger VCX1 | FUN_006476 |
| **Proton gradient** | H<sup>+</sup>-ATPase PMA1; V-ATPase subunits VMA1, VMA2, VPH1 | FUN_006808; FUN_000158, FUN_003254, FUN_001894 |
| **K<sup>+</sup> uptake** | HAK-type transporters; TRK1 | FUN_004306, FUN_009044; FUN_000163 |
| **Osmosensing (HOG)** | Sln1/TcsB, Ypd1, Ssk1, Ssk2, Pbs2, Hog1-type MAPKs | FUN_008947, FUN_002835, FUN_000740, FUN_004412, FUN_003877, FUN_004254, FUN_009420 |
| **Ionic signalling** | Calcineurin CNA1, CNB1; transcription factor CrzA | FUN_000100, FUN_004795; FUN_001184 |
| **Compatible solutes** | Glycerol synthesis (GPD1/GfdA, GPP1) and retention (Fps1, Stl1); trehalose (TPS1, TPS2); mannitol (MpdA) | FUN_006775, FUN_005837; FUN_002894, FUN_003639; FUN_003497, FUN_008328; FUN_002181 |

Phytohormone production was not assayed in this study, but candidate genes were identified for three routes (Table 3). Two genes were assigned to ACC deaminase (EC 3.5.99.7): FUN_006661, annotated as such, and FUN_000448, annotated as a pyridoxal-phosphate-dependent enzyme. For auxin, the genome encodes the complete tryptophan biosynthetic pathway (TRP2, FUN_003446; TRP4, FUN_001950; TRP3, FUN_004234; tryptophan synthase, FUN_003233, FUN_003573), an ARO8 aromatic aminotransferase (FUN_003207) for the entry step of the indole-3-pyruvate route and an aromatic L-amino acid decarboxylase (FUN_009706) of the tryptamine route; no tryptophan 2-monooxygenase was found, and the indole-3-acetonitrile route is represented only by two generic nitrilases (FUN_003565, FUN_006226). For cytokinins, a tRNA-isopentenyltransferase (FUN_004731) and two LOG-like phosphoribohydrolases (FUN_003275, FUN_004765) were identified. No bifunctional ent-copalyl diphosphate/ent-kaurene synthase was found, and no terpene BGC (Figure 2E) matched a gibberellin cluster.

Finally, the secretome includes proteins associated with root colonisation in other fungi (Table 3): three LysM proteins with signal peptides (FUN_001690, FUN_008586, FUN_009355), two of them predicted effectors; six hydrophobins, three of class I (FUN_000077, FUN_000268, FUN_000429) and two of class II (FUN_009018, FUN_009291), and secreted members of GH10 (FUN_009088) and AA9 (FUN_005814, FUN_009388). A cerato-platanin predicted as an apoplastic effector (FUN_002426), related to elicitors of induced systemic resistance (Djonović et al., 2007), was also found.

### P. melinii ‘isolate 2’ tolerates salinity and promotes Arabidopsis growth under salt stress

Beyond its growth-promoting activity, the ecology of the species raises a second question. *P. melinii* has been recovered from the rhizosphere of the halophyte grass *Aeluropus littoralis* and reported as plant growth promoting on tobacco (A8 strain; Tarroum et al., 2021), but it has not been tested for benefit to a plant under salinity. The isolate examined here, *P. melinii* ‘isolate 2’, was recovered from a habitat with no saline influence, the roots of wild *A. thaliana* in central Spain, in a Mediterranean evergreen oak woodland of *Quercus rotundifolia* and *Q. coccifera* on acidic soil developed over granite (Pagán et al., 2010; García et al., 2013), and has been described as growth-promoting in several plant species (Gutiérrez-Manso et al., 2026). We therefore tested whether the isolate itself tolerates salt and whether its growth-promoting effect extends to a host grown under salinity conditions.

Radial growth at 7 dpi did not differ from the untreated control at any concentration between 10 and 150 mM NaCl (p ≥ 0.46; Figure 5A, B). Only at 300 mM did the colony fall below the control, from 4.74 to 4.42 cm, a reduction of 0.32 cm or 6.8% (p = 0.0049). At 1 M the colony, although affected, still reached about half the control diameter, but growth was almost abolished at 3 M NaCl (Figure 5A, B). Conidiation behaved differently. Mean conidial yield was below the control at every concentration, but the reduction was statistically significant only at 300 mM, where yield fell to 0.44 times the control (p = 0.029; p ≥ 0.24 from 10 to 150 mM; Figure 5C); conidia were nearly absent at 1 and 3 M. Conidiation was therefore the more salt-sensitive trait, and the reduction at 300 mM was larger in proportional terms than the reduction in diameter at the same concentration.

**Figure 5.**
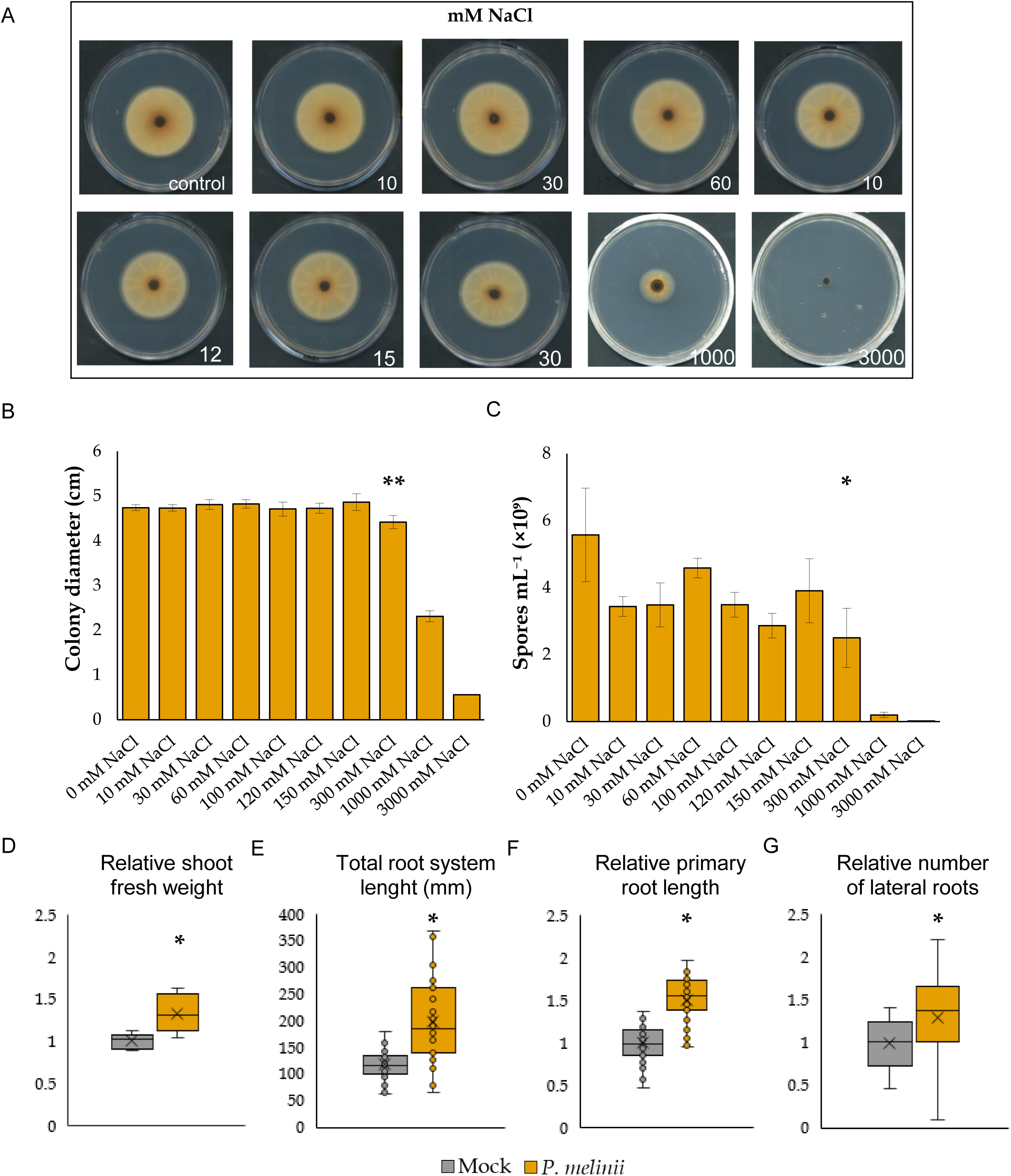
Salinity tolerance of *Penicillium melinii* ‘isolate 2’ and its effect on *Arabidopsis thaliana* under salt stress. (A) Colonies after 7 days on potato dextrose agar amended with sodium chloride at the concentrations indicated, in mM. (B) Colony diameter and (C) conidial yield at 7 dpi across the same series. Bars are means and error bars standard deviations of (3 for B, 2 for C) independent replicates, each the average of three to nine measurements; asterisks mark concentrations differing from the unamended control by a two-sided Dunnett test on a model blocking on replicate, applied to untransformed diameters in B and to log-transformed counts in C. (D–G) Effect of inoculation on *A. thaliana* grown on half-strength MS medium containing 100 mM sodium chloride: relative shoot fresh weight (D), total root system length (E), relative primary root length (F) and relative number of lateral roots (G). Grey, mock-inoculated; orange, inoculated with *P. melinii* ‘isolate 2’. Boxes span the interquartile range, the horizontal line is the median, the cross is the mean and whiskers extend to the most extreme values within 1.5 times the interquartile range; three independent experiments with 10–19 plants per treatment each; Student’s t-test, p = 0.0033 (D), p = 5.12 × 10⁻⁷ (E), p = 5.2 × 10⁻¹⁴ (F) and p = 0.0023 (G).

The genome of P. melinii ‘isolate 2’ encodes the core components of the Na⁺/K⁺ homeostasis and osmotic stress-response network (Table 3). For Na⁺ and K⁺ homeostasis, four ENA-type P-type ATPases, two of them ENA2 orthologs (FUN_004879, FUN_009227), one ENA-type (FUN_007340) and one Cta3-like (FUN_002495); a plasma-membrane Nha1-type (FUN_005033) and an endosomal NHX1-type (FUN_000670) Na⁺/H⁺ antiporter; the vacuolar cation/H⁺ exchanger VCX1 (FUN_006476); the plasma-membrane H⁺-ATPase PMA1 (FUN_006808) and the V-ATPase subunits VMA1 (FUN_000158), VMA2 (FUN_003254) and VPH1 (FUN_001894); and two HAK-type (FUN_004306, FUN_009044) and one TRK1-type (FUN_000163) K⁺ transporters. The two-component branch of the high-osmolarity glycerol (HOG) pathway is complete, comprising Sln1/TcsB (FUN_008947), Ypd1 (FUN_002835), Ssk1 (FUN_000740), Ssk2 (FUN_004412), Pbs2 (FUN_003877) and two Hog1-type MAPKs (FUN_004254, FUN_009420), as are calcineurin (CNA1, FUN_000100; CNB1, FUN_004795) and its target transcription factor CrzA (FUN_001184). Genes for glycerol synthesis (GPD1/GfdA, FUN_006775; GPP1, FUN_005837) and retention/uptake (Fps1, FUN_002894; Stl1, FUN_003639), trehalose synthesis (TPS1, FUN_003497; TPS2, FUN_008328) and mannitol synthesis (MpdA, FUN_002181) were also present.

We then asked whether this tolerance is accompanied by a benefit to a host exposed to salt. Plants were grown on half-strength Murashige and Skoog medium containing 100 mM sodium chloride, a concentration at which the isolate itself grows as well as in its absence on PDA (Figure 5A-B). Shoot fresh weight was higher in inoculated plants (Figure 5D; p = 0.0033) and the effect on the root system was larger than on the shoot: total root system length increased from 11.8 to 19.8 cm (Figure 5E; p = 5.12 × 10^-7^), primary root length also increased (Figure 5F; p = 5.2 × 10^-14^), and the number of lateral roots increased to a lesser extent (Figure 5G; p = 0.0023). ‘Isolate 2’ therefore tolerates moderate salinity and promotes the growth of *Arabidopsis* under it, although neither observation establishes the underlying mechanism.

## Discussion

*Penicillium melinii* ‘isolate 2’ was recovered from a soil without saline influence, yet its mycelial growth was unaffected by NaCl up to 150 mM and persisted at 1M, and it improved the root system of *Arabidopsis* at 100 mM NaCl. The genome provides candidate genes for the positive nutrient-mobilising traits, for hormonal routes and root colonisation and for ion and osmotic homeostasis (Table 3). We discuss first what distinguishes this genome and then how its gene content relates to the phenotypes observed.

### A compact genome at the lower end of the genus

The assembly (29.39 Mb) is smaller than the mean of 33.27 Mb obtained for 93 long-read *Penicillium* genomes (range 25.4–46.5 Mb), and its 9,566 protein-coding genes fall just below the lowest value recorded when 104 genomes were annotated together, whose predicted proteomes ranged from 9,591 to 14,319 proteins with a mean of 11,976 (Petersen et al., 2023). The short-read assembly of the same isolate yielded 10,514 predicted coding sequences (Gutiérrez-Manso et al., 2026), so the gene count depends on the assembly and annotation strategy. The low gene number does not reflect an incomplete resource, since the annotation recovered 98.7% of complete BUSCOs, comparable to the mean of 97.87% in that survey, although lineage datasets and annotation pipelines differ. Contiguity is high, with six nuclear scaffolds, two of them telomere-to-telomere, whereas the assemblies of that survey comprised five to 28 contigs. The final assembly nevertheless depends on reference-guided scaffolding, so we regard it as near-chromosome-level.

### A reduced repertoire in a lineage distinct from *P. bilaiae*

Genome-scale phylogeny places ‘isolate 2’ next to *P. maclennaniae*, with *P. citrinum* in the same broader clade and *P. bilaiae*, the species marketed as a phosphate-solubilising inoculant (Leggett et al., 2007), as sister to that broader clade. Plant growth-promoting *Penicillium* species are thus scattered across the genus, and the traits shared by ‘isolate 2’ and *P. bilaiae* cannot be attributed to close ancestry; whether they arose independently or were retained from a distant ancestor and lost elsewhere cannot be decided without screening the intervening lineages. Gene-family overlap reinforces this point: ‘isolate 2’ shares more orthogroups exclusively with *P. bilaiae* (n = 1,561) than with the phylogenetically closer *P. citrinum* (n = 528), a pattern that probably reflects the larger proteome of *P. bilaiae* (13,606 predicted proteins) rather than evolutionary affinity.

Relative to both congeners, ‘isolate 2’ encodes the fewest CAZymes (207, 2.2% of the proteome, against 535 in *P. bilaiae* and 443 in *P. citrinum*), biosynthetic gene clusters (37 against 78 and 44), candidate effectors (109 against 206 and 141) and species-specific orthogroups (30 against 246 and 60). The contrast is sharpest with *P. bilaiae*, which also has the largest secretome, whereas ‘isolate 2’ and *P. citrinum* predict similar numbers of extracellular proteins (471 and 459) and differ mainly in effectors and CAZymes. Among CAZymes, GH18 chitinases show the strongest contraction (3 genes, against 26 in *P. bilaiae* and 19 in *P. citrinum*). Annotation coverage was highest for ‘isolate 2’, so poorer functional annotation cannot explain the smaller counts. The congener proteomes, however, derive from different gene-calling pipelines, and a conservative, evidence-driven prediction would both raise annotation coverage and lower the number of lineage-specific genes (Weisman et al., 2022), so absolute numbers should be compared with caution.

Laccase-like multicopper oxidases are scarce (two AA1 genes, one of them AA1_3) and several families acting on plant cell-wall polysaccharides (AA3_1, GH43_30, CBM67) are absent, which agrees with the negative laccase and cellulase assays and points to a lifestyle not based on lignocellulose degradation. The Biolog profile is consistent with this picture: growth was strongest on di- and oligosaccharides such as maltose, cellobiose and gentiobiose, whereas no cellulase halo formed on CMC, and growth on dextrin (5.0-fold) and glycogen (3.7-fold) shows that α-glucans are used even though no halo formed on starch. Repertoire size alone, however, does not define lifestyle (Mesny et al., 2021). ‘Isolate 2’ retains secreted members of GH10 and AA9, two families associated with endophytism in that study, and lacks the pectate lyase subfamily PL1_7, which was linked to aggressive colonisation and is present in both congeners. A reduced degradative and effector repertoire is therefore compatible with, though not evidence for, a beneficial root association, which in this isolate rests on the phenotype (Gutiérrez-Manso et al., 2026).

### Nutrient mobilisation: genome and plate assays largely agree

Each positive assay of nutrient mobilisation except zinc solubilisation (siderophores, phosphate, zinc, urease, ammonia and protease) has a genomic counterpart (Table 3). Siderophore production is matched by a hydroxamate pathway whose committed enzyme, SidA, is essential for siderophore biosynthesis in *Aspergillus fumigatus* (Schrettl et al., 2004) and lies next to an NRPS, and by a second locus with genes similar to *sidH*, *sidF* and *sidD*, which are required for extracellular fusarinine-type siderophores (Schrettl et al., 2007; Yasmin et al., 2012). This makes the trait potentially relevant to the plant, since fusarinines and dimerum acid from *P. chrysogenum* improved iron utilisation by both cucumber and maize (Hördt et al., 2000).

For phosphate solubilisation, three citrate synthases and the glyoxylate-shunt enzymes isocitrate lyases and malate synthase are consistent with the organic acid mechanism characterised in *P. bilaiae* (Cunningham and Kuiack, 1992), although these enzymes belong to central metabolism and are broadly present in fungal genomes. Zinc solubilisation, for which no specific gene was identified, is most simply explained by the same acids. That halos formed from starting pH values of 5.7, 7.0 and 8.0 should not, however, be read as solubilisation at alkaline pH. Pikovskaya medium is unbuffered, and two phosphate-solubilising bacteria active in unbuffered media failed to solubilise rock phosphate once the medium was buffered (Gyaneshwar et al., 1998), whereas P. bilaiae solubilised more rock phosphate in buffered (pH 7) than in non-buffered medium, by increasing citric and oxalic acid production (Takeda and Knight, 2006). Repeating the assay in buffered medium and recording the final pH is therefore necessary before extending the trait to alkaline saline soils.

Organic phosphorus, which none of the assays tested, is a separate target for eight secreted phosphatases: three histidine acid phosphatases including PHO5, three PhoA-type acid phosphatases, a purple acid phosphatase and an alkaline phosphatase (Table 3). Finally, the urease URE1 and 16 secreted proteases, among them the subtilisin PRB1, the aspartic proteases PepA and an M35 metalloprotease (Table 3), are consistent with the urease activity, gelatine hydrolysis and ammoniagelatine release from peptone. These capacities could contribute to the growth-promoting activity of ‘isolate 2’ in soil, although none has yet been linked to the plant phenotype.

### Candidate determinants of the root response

Three hormonal mechanisms that could contribute to growth promotion stood out from the genome analysis. The first is ACC deaminase, with one gene annotated as such and second candidate annotated only as a pyridoxal-phosphate-dependent enzyme (Table 3). The enzyme has been purified from *Penicillium citrinum* (Jia et al., 1999), and its silencing in *Trichoderma asperellum* reduced the promotion of root elongation (Viterbo et al., 2010). By removing the ethylene precursor, it could relieve ethylene-mediated inhibition of root growth under salt, but the activity must be demonstrated. The second is the indole-3-pyruvate route of auxin synthesis, established genetically in *Ustilago maydis* (Reineke et al., 2008) and implicated by inhibitor treatment in IAA production by *P. citrinum* B9 (Chen et al., 2023). ‘Isolate 2’ encodes the complete tryptophan pathway and the ARO8 aminotransferase that performs the entry step (Table 3), although these genes are widely conserved in fungi. Both *P. melinii* A8 (Tarroum et al., 2021) and ‘isolate 2’ (Gutiérrez-Manso et al., 2026) release Salkowski-reactive indolic compounds, increasingly with tryptophan, although this assay is not specific for IAA. The third, the tRNA-degradation route to cytokinins, comprises a tRNA-isopentenyltransferase and two LOG-like phosphoribohydrolases (Table 3); its first enzyme is conserved in most fungi (Chanclud et al., 2016) and therefore does not distinguish this isolate either. The canonical gibberellin pathway is absent, which removes one explanation without excluding gibberellin production by another route.

Beyond hormones, the secretome contains candidates for the root interface (Table 3). A cerato-platanin predicted as an apoplastic effector is related to Sm1, which *Trichoderma virens* requires to induce systemic resistance in maize but not to colonise its roots (Djonović et al., 2007). Three secreted LysM proteins, two of them predicted effectors, belong to a family that in beneficial fungi protects hyphae from chitinases and dampens chitin-triggered defences: silencing *RiSLM* greatly reduced the colonisation of *Medicago truncatula* roots by *Rhizophagus irregularis* (Zeng et al., 2020), and Tal6 of *Trichoderma atroviride* modulates the activation of plant defences (Romero-Contreras et al., 2019). Hydrophobins are a further candidate, since deletion of the class I-like hydrophobin gene *TasHyd1* severely impaired root attachment and colonisation by *Trichoderma asperellum* (Viterbo and Chet, 2006). Whether any of these proteins is expressed during colonisation of *Arabidopsis* remains to be tested.

### Salt tolerance without a saline origin

The genome offers candidates for the salt tolerance of the mycelium (Table 3): four P-type ATPases of the ENA family (two ENA2, one ENA-type and one Cta3-like), the Na⁺/H⁺ antiporters Nha1 and NHX1, two HAK and one TRK K⁺ transporters, and genes for glycerol synthesis (GPD1, GPP1), and retention (Fps1), and uptake (Stl1) . Osmotic signalling is covered by the two-component branch of the HOG pathway, from Sln1 through Ypd1, Ssk1, Ssk2 and Pbs2 to two Hog1-type kinases. In *Aspergillus nidulans* this is the only branch that activates the osmotic response, although it depends on the response regulator SskA rather than on the Sln1 orthologue TcsB (Furukawa et al., 2005). Most of these genes are conserved across filamentous ascomycetes, so their presence does not by itself explain the tolerance of this isolate, and they bear on how the fungus persists under salt rather than on how it helps the plant. The exception is the ENA family, whose homologues in *Serendipita* indica are expressed during co-cultivation with *Arabidopsis* under salinity (Lanza et al., 2019; Haro et al., 2023). ENA ATPases also sustain growth at alkaline pH in *A. nidulans* and are induced at high pH in *S. indica* (Lanza et al., 2019; Markina-Iñarrairaegui et al., 2020), which makes them relevant to alkaline saline soils; whether the ENA genes of ‘isolate 2’ act in either way is a testable hypothesis.

*Penicillium melinii* A8, isolated from the rhizosphere of the halophyte *Aeluropus littoralis* in a saline sebkha, also grew at up to 1 M NaCl (Tarroum et al., 2021). ‘Isolate 2’, an endophyte from roots of Arabidopsis growing on an acidic granitic soil (García et al., 2013), showed no cost to radial growth up to 150 mM and still grew at 1 M, at about half the control diameter. The two responses differ in shape rather than simply in degree, and they cannot be compared directly, because incubation time and concentration range differed between the assays. They nevertheless indicate that a saline origin is not a reliable predictor of tolerance to moderate salinity, since A8 was affected from 100 mM, and that this tolerance in ‘isolate 2’ did not require selection in a saline habitat. Isolates from non-saline soils therefore merit evaluation as inoculants for saline ones. Moreover, A8 was tested on tobacco without salt, so the present work provides, to our knowledge, the first evidence of plant growth promotion under salinity for this species.

## Author Contributions

Conceptualization: SGS, LGM, JC; Data curation: SGS, LGM; Formal analysis: SGS, LGM; Funding acquisition: LU, JN, AVM, SS, EC, JCP, JC; Investigation: SGS, LGM, IDA, FB, LBC, CC, LU, JN, AVM, SS, EC, JCP, JC; Methodology: SGS, LGM, IDA, FB, LBC, CC, LU, JN, AVM, SS, EC, JCP, JC; Resources: LU, JN, AVM, SS, EC, JCP, JC; Supervision: SGS, CC, LU, JN, AVM, SS, EC, JCP, JC; Validation: SGS, LGM, IDA, FB; Visualization: SGS, LGM, FB, JC; Writing – original draft: SGS, LGM, JC; Writing – review & editing: SGS, LGM, IDA, FB, LBC, CC, LU, JN, AVM, SS, EC, JCP, JC.

## Generative AI Statement

The authors declare that generative AI was used in the preparation of this manuscript. A large language model (Claude Opus 5.5, Anthropic) was used for two purposes. First, it was used to query the functional annotation tables generated in this study and to identify genes that could act as candidates for plant growth-promoting and salt-tolerance functions, as described above. The same model was used to locate published studies on functionally characterized fungal orthologs. Second, it assisted with manuscript writing: drafting, correcting the English, linking ideas between sections and removing redundancies. All candidate genes, references and text produced or edited with the model were checked by the authors against the original annotation data and the published literature. The authors take full responsibility for the content of this article.

## Supplementary Material

**Supplementary Figure 1.** Circular map of the mitochondrial genome of *Penicillium melinii* ‘isolate 2’ (26,730 bp). Features are colour-coded by category: protein-coding genes (pink), tRNA genes (dark orange) and rRNA genes (gold). Gene names are displayed outward; tick marks indicate position in kilobases (kb). The linear start (position 1) was set by Circlator fixstart at the ATG start codon of a Prodigal-predicted open reading frame; cox1 spans the resulting join point. The gene complement is complete and consistent with a *Penicillium* sp. mitochondrion, comprising the three cytochrome oxidase subunits (cox1–cox3), apocytochrome b (cob), three ATP synthase subunits (atp6, atp8, atp9), seven NADH dehydrogenase subunits (nad1–nad6 and nad4L), the small subunit ribosomal protein rps3, the large and small subunit ribosomal RNAs (rrnL, rrnS), and the full transfer RNA complement. The map was generated with a custom Python script.

**Supplementary Figure 2.** Maximum-likelihood phylogenetic tree of *P. melinii* ‘isolate 2’ and 16 reference taxa (15 *Penicillium* spp. + *Aspergillus fumigatus* as outgroup), inferred with FastTree v2.2.0 from a concatenated supermatrix of 2,223,900 amino acid positions derived from single-copy BUSCO orthologues. Node circles indicate SH-like local support values (colour scale, 0–1.0); scale bar = 0.02 substitutions per site. *P. melinii* ‘isolate 2’ (highlighted) is placed sister to *P. maclennaniae* (SH = 1.0), with *P. atrosanguineum* and *P. corylophilum* as successive sister lineages.

**Supplementary Tables 1-5.**

