## Supplementary Figure for "Genomic and phenotypic characterisation of *Penicillium melinii*, a plant growth-promoting root endophyte that remains effective under salt stress"

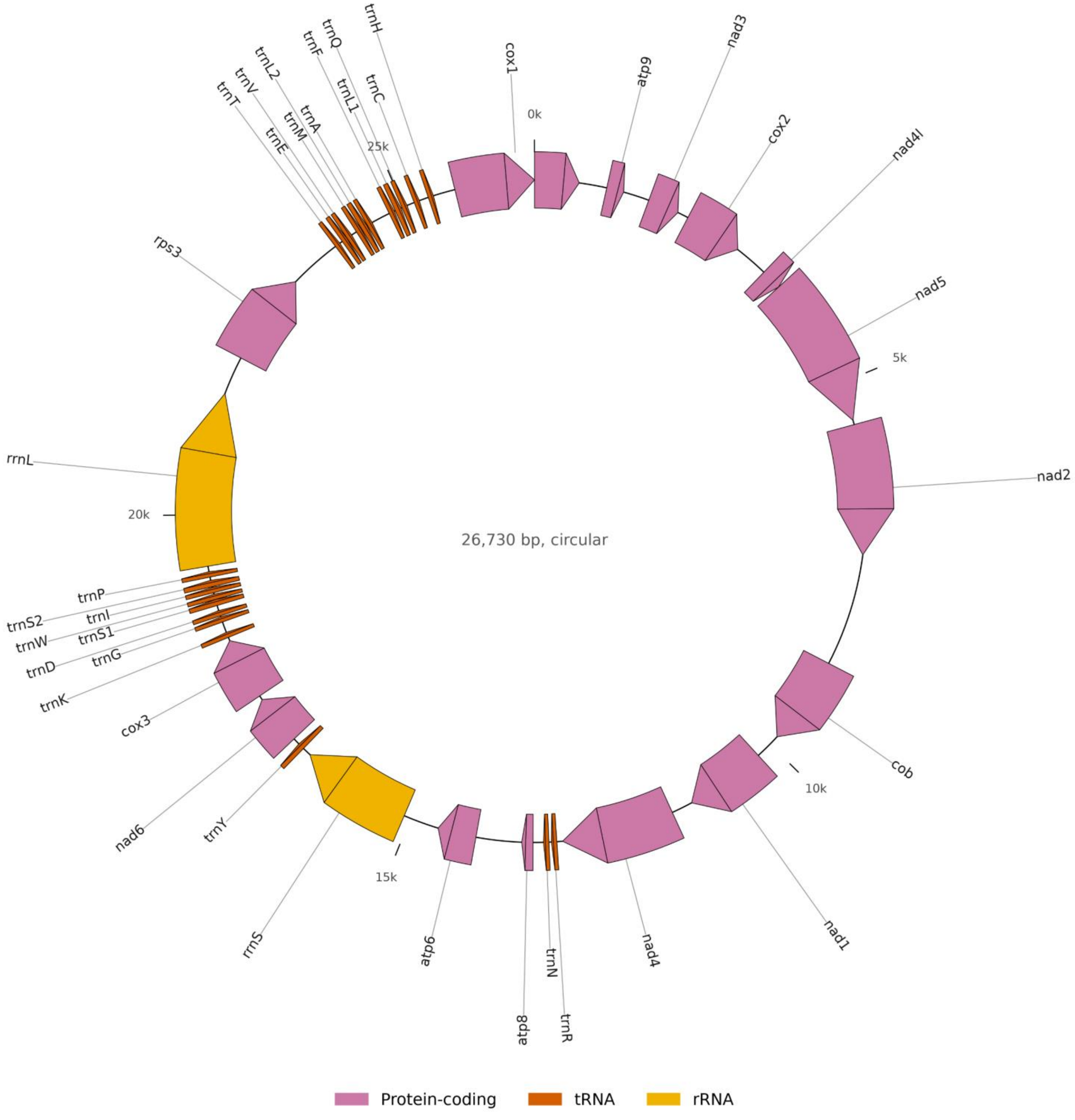


**Supplementary Figure 1.** The mitochondrial genome of *Penicillium melinii* 'isolate 2'. The molecule assembled as a single circular sequence of 26,730 bp. Pink, protein-coding genes; orange, transfer RNA genes; yellow, ribosomal RNA genes; the scale is in kb from the arbitrary origin. The complement is that expected of a *Penicillium* sp. mitochondrion and is complete: the three cytochrome oxidase subunits (*cox1-cox3*), apocytochrome b (*cob*), three ATP synthase subunits (*atp6, atp8, atp9*), seven NADH dehydrogenase subunits (*nad1-nad6* and *nad4L*), the small subunit ribosomal protein *rps3*, the large and small subunit ribosomal RNAs (*rrnL, rrnS*), and the transfer RNAs.

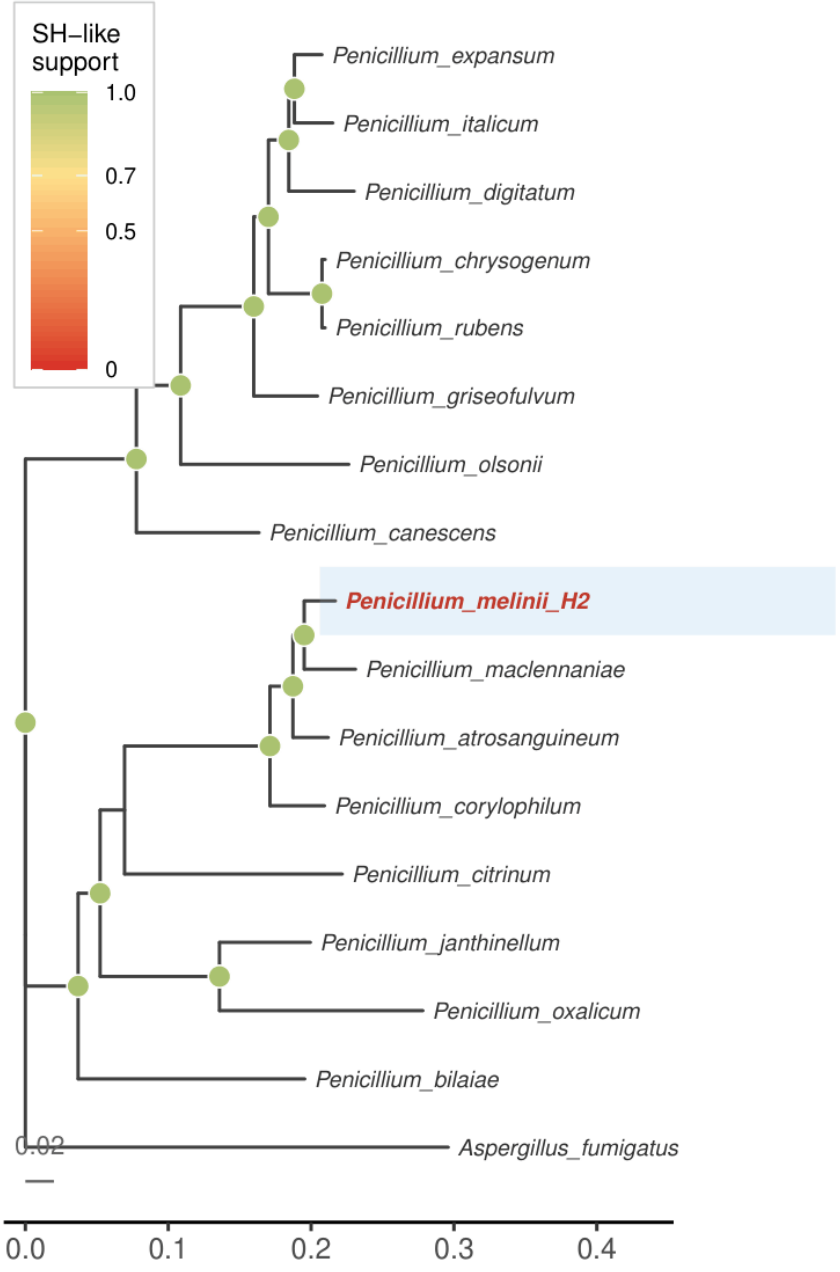


**Supplementary Figure 2.** Maximum-likelihood phylogenetic tree of *P. melinii* 'isolate 2' and 16 reference taxa (15 *Penicillium* spp. + *Aspergillus fumigatus* as outgroup), inferred with FastTree v2.2.0 from a concatenated supermatrix of 2,223,900 amino acid positions derived from single-copy BUSCO orthologues. Node circles indicate SH-like local support values (colour scale, 0–1.0); scale bar = 0.02 substitutions per site. *P. melinii* 'isolate 2' (highlighted) is placed sister to a clade comprising *P. maclennaniae*, *P. atrosanguineum*, and *P. corylophilum*, with maximum node support.
